# Prototype mRNA vaccines imprint broadly neutralizing human serum antibodies after Omicron variant-matched boosting

**DOI:** 10.1101/2024.01.03.574018

**Authors:** Chieh-Yu Liang, Saravanan Raju, Zhuoming Liu, Yuhao Li, Guha A. Arunkumar, James Brett Case, Seth J. Zost, Cory M. Acreman, Deborah Carolina Carvalho dos Anjos, Jason S. McLellan, James E. Crowe, Sean P.J. Whelan, Sayda M. Elbashir, Darin K. Edwards, Michael S. Diamond

## Abstract

Immune imprinting is a phenomenon in which an individual’s prior antigenic experiences influence responses to subsequent infection or vaccination. Here, using antibody depletion and multiplexed spike-binding assays, we characterized the type-specificity and cross-reactivity of serum antibody responses after mRNA vaccination in mice and human clinical trial participants. In mice, a single priming dose of a preclinical version of mRNA-1273 vaccine encoding Wuhan-1 spike minimally imprinted serum responses elicited by Omicron boosters, enabling a robust generation of type-specific antibodies. However, substantial imprinting was observed in mice receiving an Omicron booster after two priming doses of mRNA-1273, an effect that was mitigated by a second booster dose of Omicron mRNA vaccine. In humans who received two BA.5 or XBB.1.5 Omicron-matched boosters after two or more doses of the prototype mRNA-1273 vaccine, spike-binding and neutralizing serum antibodies cross-reacted with circulating Omicron variants as well as more distantly related sarbecoviruses. Because the serum neutralizing response against Omicron strains and other sarbecoviruses was completely abrogated after pre-clearing with the Wuhan-1 spike protein, antibodies induced by XBB.1.5 boosting in humans focus on conserved epitopes shaped and shared by the antecedent mRNA-1273 primary series. Our depletion analysis also identified cross-reactive neutralizing antibodies that recognize distinct epitopes in the receptor binding domain (RBD) and S2 proteins with differential inhibitory effects on members of the sarbecovirus subgenus. Thus, although the serum antibody response to Omicron-based boosters in humans is dominantly imprinted by prior immunizations with prototype mRNA-1273 vaccines, this outcome can be beneficial as it drives expansion of multiple classes of cross-neutralizing antibodies that inhibit infection of emerging SARS-CoV-2 variants and extend activity to distantly related sarbecoviruses.

## INTRODUCTION

Multiple vaccines targeting the spike protein of the Wuhan-1 isolate of Severe acute respiratory syndrome coronavirus 2 (SARS-CoV-2) were rapidly developed to control the COVID-19 pandemic. Despite the initial success of these vaccines in limiting symptomatic SARS-CoV-2 infections, the evolution of variant strains with increasing numbers of escape mutations in the spike protein has limited their efficacy^1–4^. Indeed, decreased serum neutralization and breakthrough infections with the Omicron BA.1 and subsequent lineage variants have been reported in vaccinated individuals^5–7^. In response to this loss of protection, vaccine boosters encoding variant spike proteins were designed and deployed. Immunization with vaccine boosters encoding the spike protein of B.1.351 or BA.1 enhanced neutralizing antibody responses against both the ancestral virus and the corresponding variant^2,8–11^. As comparable results were achieved with the original Wuhan-1 spike-based vaccine boosters^2,11^, it remains unclear how antibody responses differ after boosting with variant matched or non-matched spike proteins.

Although the goal of variant-matched boosters is to enhance protection against the variant strains, this outcome could be limited by effects of prior SARS-CoV-2 infection or vaccination. Immune imprinting describes the phenomenon in which immunity against a second antigen is shaped by antecedent exposures to a prior, related antigen, and is thought to be mediated by the recall of cross-reactive memory B cells (MBCs) at the expense of stimulating *de novo* B cell responses. The phenomenon, first termed “original antigenic sin”, described the observation that individuals had higher antibody titers against influenza viruses that they had been exposed to during childhood instead of more contemporary strains^12^.

Multiple studies have suggested imprinting effects by prior SARS-CoV-2 antigen exposures on subsequent infections with variant strains or immunizations with variant-matched vaccines^13–16^. For example, in one cohort of triply-vaccinated (BioNTech BNT162b2) healthcare workers, BA.1 infection increased BA.1-reactive antibody responses in the serum of infection-naïve individuals but not in those who had also experienced Wuhan-1 infections^13^. Studies in animals and humans have shown relatively small differences in variant spike-binding antibody titers between subjects boosted with Wuhan-1-encoding vaccines and variant-matched vaccines^2,9–11,17^. Imprinting effects on SARS-CoV-2 immunity generally have been studied by analyzing the spike-binding reactivity of B cells via flow cytometry or through B cell repertoire and monoclonal antibody (mAb) analysis^9,11,14,16^. Rhesus macaques immunized with a primary series of the Wuhan-1 spike-encoding mRNA-1273 vaccine and boosted heterologously with B.1.351- or BA.1-matched vaccines showed poor variant-specific MBC responses^9,11^. The vast majority of mAbs derived from MBCs isolated from human mRNA vaccine recipients primed with the Wuhan-1 spike-encoding vaccine and boosted with the BA.1-matched vaccine were cross-reactive to the Wuhan-1 strain, as very few BA.1-specific mAbs (< 1%) were detected^16^. Although these results indicate a strong imprinting effect in the MBC compartment, the serum antibody profile may differ since it reflects contributions from the antibody-secreting cells, including long-lived plasma cells (LLPCs) in the bone marrow, which may not share the same antigen specificities as MBCs^18^. A key question remains as to whether heterologous booster vaccines induce serum antibody responses that are type-specific to the variant strain or cross-reactive to the original priming strain, and what fraction each of these contributes to neutralization of the variant strain.

Here, we determined the presence and function of virus-specific and cross-reactive antibodies in the serum of heterologously vaccinated mice and human clinical trial participants using antibody depletion, multiplexed spike-binding detection, and neutralization assays. We evaluated the imprinting effect of antecedent mRNA-1273 vaccines encoding Wuhan-1 spike on mouse and human serum antibody responses after boosting with one or two doses of mRNA vaccines encoding Omicron BA.1, BA.5, or XBB.1.5 spike proteins.

## RESULTS

### Multiplexed quantification of spike-reactive antibodies

To measure the reactivity of serum antibodies against multiple spike protein variants of SARS-CoV-2, we developed a flow cytometry-based multiplexed assay. Biotinylated spike proteins of different SARS-CoV-2 strains were bound to streptavidin-coated microparticles with varying intensities of SPHERO™ yellow fluorophore, incubated with serum samples, and the level of antibody binding to each spike protein was determined by flow cytometry (**Fig. 1a** and **Extended Data Fig. 1a**). To normalize antibody titers acquired across experiments, a mAb, CV3-25, that recognizes a highly conserved epitope in the S2 stem region^19^ was run in parallel to create a standard curve (**Fig. 1a-b**).

**Figure 1.**
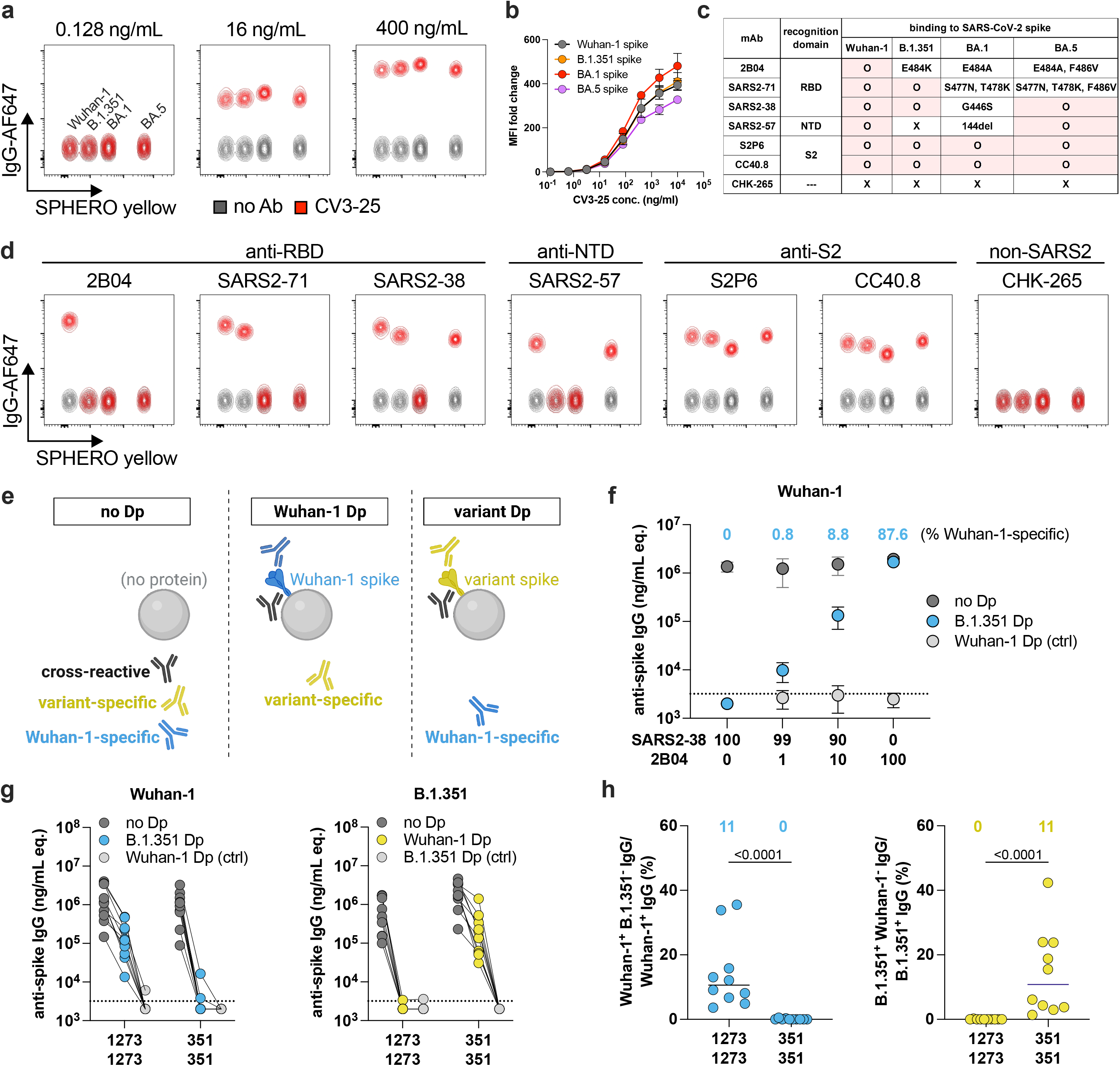
A multiplexed spike-binding assay and antibody depletion assay. **a,** Representative flow cytometry plots of mAb CV3-25 binding to Wuhan-1, B.1.351, BA.1, and BA.5 spike-charged detection beads at the indicated antibody concentrations. **b**, A CV3-25 standard curve of binding to Wuhan-1, B.1.351, BA.1, and BA.5 spike-charged detection beads (pooled from two experiments). **c-d**, SARS-CoV-2 spike recognition and binding characteristics of the indicated mAbs (**c**) and flow cytometry plots (**d**) of mAb binding to Wuhan-1, B.1.351, BA.1, and BA.5 spike-charged detection beads (three experiments). A mAb that binds chikungunya virus (CHK-265) was included as negative control. **e**, Schematic of the anti-spike antibody depletion assay. **f**, Wuhan-1 spike-reactive antibody titers of mAbs SARS2-38 and 2B04 mixed at the indicated ratios and pre-cleared with empty depletion beads (no Dp), B.1.351 spike-loaded beads (B.1.351 Dp), or Wuhan-1 spike-loaded beads (Wuhan-1 Dp) (numbers at the top indicate the derived fraction of Wuhan-1 spike-specific antibody; three experiments; dotted lines show the limit of detection, LOD). **g**, Antibody binding of 3-week post-boost sera harvested from mice primed with 0.25 μg and boosted one month later with 1 μg of mRNA-1273 (1273/1273) or mRNA-1273.351 (351/351) (n = 10, two experiments) to Wuhan-1 and B.1.351 spike proteins as measured by the multiplexed spike-binding assay. Sera were pre-cleared with empty beads, B.1.351 spike-loaded beads, or Wuhan-1 spike-loaded beads (connecting lines represent sera from the same mouse; dotted lines show the LOD; values at the LOD are plotted slightly below the LOD for visualization). **h**, Percentages of Wuhan-1 (*left*) and B.1.351 (*right*) spike-specific antibodies, derived from **g**. The fraction of Wuhan-1 spike-specific IgG is calculated by dividing the Wuhan-1 spike-binding titer of B.1.351 spike depleted sera (*blue circles in **g** left panel*) by that without depletion (*dark grey circles in **g** left panel*) from the same individual (B.1.351 Dp/no Dp) (type-specific titers at the LOD in **g** are designated 0% type-specific response in **h**; numbers at the top illustrate median values). **h**, Mann-Whitney test with Bonferroni correction.

To validate the assay, a panel of mAbs with SARS-CoV-2 spike protein reactivity across the receptor-binding domain (RBD), N-terminal domain (NTD), and S2 stem region were tested (**Fig. 1c-d**). 2B04, a mAb generated from mice primed and boosted with Wuhan-1 spike proteins, bound Wuhan-1 spike but not the other variant spike proteins tested, consistent with its loss-of-binding to variants with mutations at residue 484^20,21^. SARS2-71 bound to Wuhan-1 and B.1.351 but not to BA.1 and BA.5 spike proteins, consistent with this mAb losing binding to spike proteins with S477N mutations^22^. MAb SARS2-38 bound to Wuhan-1, B.1.351 and BA.5 but not BA.1 spike, which harbors a mutation at residue 446, a key contact site of this mAb^22^. NTD-binding mAb SARS2-57 bound to Wuhan-1 and BA.5 but not B.1.351 and BA.1 spike proteins, presumably due to the deletions at residues 242 to 244 and 144, respectively^23^. In comparison, the S2-binding mAbs S2P6 and CC40.8^24,25^ bound to all spike proteins tested. Together, the binding characteristics of the tested mAbs support the utility of the multiplexed spike-binding assay in distinguishing the spike-binding reactivity of antibodies. As an additional validation step, we showed that the anti-spike titers of the mAbs obtained from the multiplexed spike-binding assay correlated with endpoint titers achieved by ELISA (**Extended Data Fig. 1b**).

### Measurement of virus type-specific antibodies in serum

Although the multiplexed assay can measure antibody binding titers to multiple spike proteins, it cannot discern the fraction of virus-specific and cross-reactive antibodies. Accordingly, we designed a depletion assay to distinguish variant type-specific and cross-reactive antibody responses using streptavidin-coated magnetic beads loaded with a biotinylated spike protein of interest (**Fig. 1e**). Binding analysis of the pre-cleared flow through sera (**Fig. 1f-h**) provides quantitative information on the antibodies that are non-reactive to the spike protein on the depletion beads but reactive to the spike proteins on the detection beads. By coupling the assays, we can measure serum antibodies that bind the spike protein of one SARS-CoV-2 strain but not another.

To validate this assay, we first tested SARS-CoV-2 Wuhan-1 strain-specific (2B04) or Wuhan-1 and B.1.351 cross-reactive (SARS2-38) mAbs using our workflow. As expected, the binding of SARS2-38 to Wuhan-1 spike-charged beads was abolished when this cross-reactive mAb was pre-cleared with either Wuhan-1 or B.1.351 spike proteins (**Fig. 1f** and **Extended Data Fig. 1c**). In comparison, the binding to Wuhan-1 spike-coated beads of mAb 2B04, which does not react with B.1.351 spike protein, was minimally impacted by pre-clearing with B.1.351 spike-coated beads. To verify the ability of the assay to distinguish samples containing mixtures of antibodies of more than one specificity, SARS2-38 and 2B04 were mixed at 99:1 and 90:10 ratios. Depletion with B.1.351 spike protein decreased the Wuhan-1 spike binding titer by 99.2% and 91.2%, respectively, consistent with the expectation that 99% or 90% of the antibody mixture cross-reacts with the B.1.351 spike protein. Thus, our assays can detect the fraction of type-specific antibodies within a mixture with reasonable dynamic range.

To evaluate virus type-specific antibodies in sera using the depletion and multiplexed binding assays, samples from preclinical versions of mRNA-1273 (encodes Wuhan-1 spike) or mRNA-1273.351 (encodes B.1.351 spike) vaccine primed-boosted mice (1273/1273 and 351/351, respectively) were depleted with Wuhan-1 or B.1.351 spike proteins and then measured for binding reactivity. Complete depleting activity by Wuhan-1 and B.1.351 spike proteins was confirmed by loss of serum antibody binding to Wuhan-1 and B.1.351 spike-coated detection beads, respectively (**Fig. 1g**, *light grey circles*). To determine whether the mRNA-1273 vaccine induced Wuhan-1 spike-specific antibodies, 1273/1273 sera were incubated first with B.1.351 spike-coated depletion beads, and then the pre-cleared sera were evaluated for residual binding to Wuhan-1 spike detection beads. Depletion of 1273/1273 sera with B.1.351 spike proteins led to reduced Wuhan-1 spike-binding reactivity, with only a fraction (approximately 11%) of the Wuhan-1 spike-reactive response not cross-reacting with B.1.351 spike (**Fig. 1g-h**, *left panels*, 1273/1273 group). As expected, nearly all Wuhan-1-reactive antibodies in 351/351-immunized mice cross-reacted with B.1.351 spike (**Fig. 1g-h**, *left panels*, 351/351 group). Similarly, B.1.351 spike-specific antibodies were detected in 351/351- but not in 1273/1273-immunized mice. Thus, our coupled depletion and binding assays distinguish type-specific from cross-reactive antibodies in serum samples.

### Variant-specific antibodies in serum of mice receiving booster vaccines encoding variant spike proteins

To begin to address whether boosting with variant-matched vaccines elicits variant-specific serum antibodies, we examined serum collected from mice that received heterologous mRNA vaccine booster injections. We immunized three cohorts of mice with either one prime/one boost, two primes/one boost, or two primes/two boosts of mRNA vaccines. Groups of 6-week-old female C57BL/6J mice were intramuscularly immunized with preclinical versions of mRNA-1273 vaccine or mRNA-1273.529 vaccine, the latter of which encodes the BA.1 Omicron spike. In the one prime/one boost cohort, mice were vaccinated one month apart with homologous (1273/1273 and 529/529) or heterologous (1273/529 and 529/1273) mRNA vaccines. Serum was collected 3 weeks after the second vaccine dose, and IgG responses to Wuhan-1 and BA.1 spike proteins were determined (**Fig. 2a**). To measure BA.1 spike-specific antibody in 1273/529-immunized mice, sera were depleted with Wuhan-1 spike and assayed for binding to BA.1 spike-coated detection beads. Despite some variation, a fraction (0 to 56%; median 10%) of BA.1-reactive IgG from 1273/529-immunized mice was virus-type-specific and did not cross-react with Wuhan-1 spike (**Fig. 2b-c**, *right panels*). Reciprocally, in the 529/1273-vaccinated group, a Wuhan-1 spike-specific IgG response was observed (5 to 17%; median 9%; **Fig. 2b-c**, *left panels*). The response to heterologous boosters was minimally imprinted by the priming dose, as statistical differences were not detected in the fraction of Wuhan-1 spike-specific IgG between 1273/1273 (range 13 to 91%; median 36%) and 529/1273 vaccinated mice (p = 0.31), nor in the fraction of BA.1 spike-specific IgG between 529/529 (range 0 to 55%; median 16%) and 1273/529 immunized mice (p > 0.99). Thus, in the context of a one prime/one boost immunization series in mice, boosting with a heterologous mRNA vaccine enabled generation of type-specific antibodies in serum, although most of the post-boost response consisted of cross-reactive antibodies.

**Figure 2.**
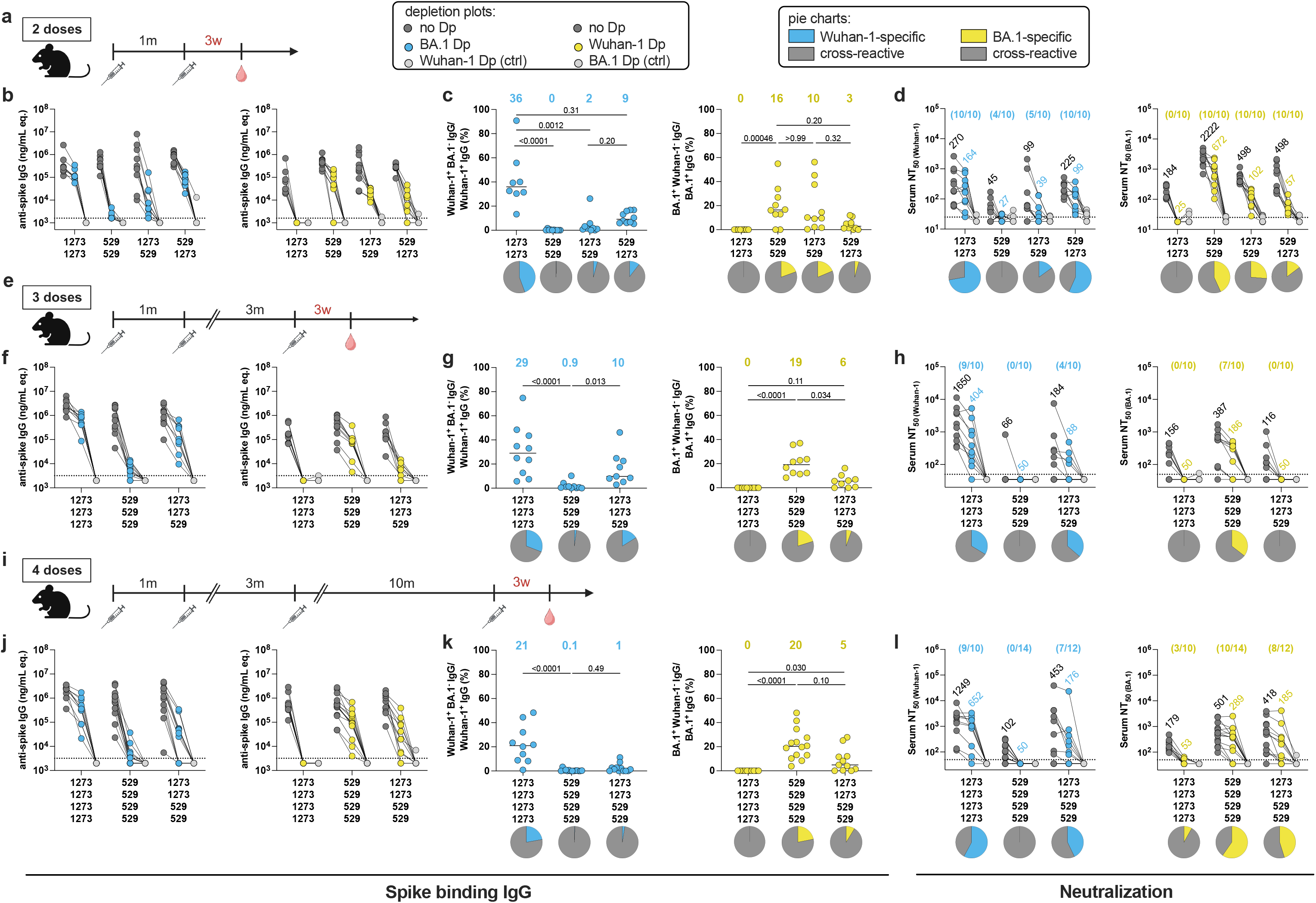
Effects of imprinting on serum binding and neutralizing antibody responses after a second dose of BA.1-matched vaccine booster in mice. **a, e, i**, Schemes of the one prime/one boost (**a**), two primes/one boost (**e**), and two primes/two boosts (**i**) immunization regimens and blood collections. **a**, Six-week-old female C57BL/6J mice were immunized twice over a 4-week interval with the indicated mRNA vaccines in **b**-**d** (n = 10, two experiments). **e**, Mice were immunized one month apart and boosted with a third dose of mRNA vaccine three months later (n = 10, two experiments). **i**, Mice were vaccinated one month apart, boosted with a third dose three months later, and received a fourth dose of mRNA vaccine ten months later (n = 10∼14, two experiments). Some of the animals in **i-l** are those described in **e-h** and boosted. All vaccines were given at 0.25 μg per dose. Sera were collected 3 weeks after the last dose. 1273, mRNA-1273; 529, mRNA-1273.529. **b, f, j**, Antibody binding of sera pre-cleared with empty beads (no Dp), BA.1 spike-loaded beads (BA.1 Dp), or Wuhan-1 spike-loaded beads (Wuhan-1 Dp) to Wuhan-1 (*left*) and BA.1 (*right*) spike proteins (connecting lines represent sera from the same mouse; dotted lines show the LOD; values at the LOD are plotted slightly below the LOD for visualization). **c, g, k**, Percentages of Wuhan-1 (*left*) and BA.1 (*right*) spike-specific antibodies. Percentages were calculated as described in Fig. 1h (numbers at the top illustrate median values). Pie charts illustrate the fraction of type-specific and cross-reactive IgG in each group. **d, h, l**, Neutralizing activity of pre-cleared sera against VSV pseudoviruses displaying Wuhan-1 (*left*) or BA.1 (*right*) spike proteins (numbers above data points indicate the geometric mean titer (GMT); fractions in parentheses at the top indicate the numbers of mice with detectable neutralization by type-specific antibodies). Pie charts illustrate the relative fractions of neutralization mediated by type-specific and cross-reactive antibodies in each group. Serum samples with an NT_50_ below the LOD after pre-clearing with empty beads (no DP) were not used in pie chart analyses. **c, g, k**, Kruskal-Wallis ANOVA with Dunn’s post-test.

To address how virus-type-specific or cross-reactive antibodies contribute to neutralization, we tested pre-cleared sera for their ability to inhibit infection of vesicular stomatitis virus (VSV)-based chimeric SARS-CoV-2 pseudoviruses^26^ (**Fig. 2d**). In mice immunized with 1273/529, BA.1-specific antibodies neutralized BA.1 pseudovirus and accounted for 20% of the neutralizing activity (**Fig. 2d**, *right panel*, NT_50_ of 102/498). In 529/1273-immunized mice, Wuhan-1-specific antibodies accounted for 44% of the neutralization of Wuhan-1 pseudovirus (**Fig. 2d**, *left panel*, NT_50_ of 99/225). Thus, in the context of a one prime/one boost vaccination scheme in mice, virus-specific antibodies elicited by heterologous booster vaccines can mediate virus neutralization.

In the two primes/one boost vaccination regimen, mice were immunized with a primary two-dose series of the preclinical version of mRNA-1273 one month apart and boosted three months later with a third dose of mRNA-1273 (1273/1273/1273) or mRNA-1273.529 (1273/1273/529) (**Fig. 2e**). Mice receiving homologous mRNA-1273.529 vaccines (529/529/529) were included for comparison. At 3 weeks after the third vaccination, BA.1-specific antibodies were detected in 78% of 1273/1273/529-vaccinated mice but at low levels (**Fig. 2f**, *right panel*), and the fraction of BA.1-specific response was lower in 1273/1273/529-vaccinated mice than in 529/529/529 mice (6% versus 19%, p = 0.034; **Fig. 2g**, *right panel*), reflecting imprinting by the primary immunization series. Although neutralization of BA.1 pseudoviruses was observed in all groups, BA.1-specific IgG in 1273/1273/529 mice did not contribute to this inhibition (**Fig. 2h**, *right panel*). At 15 weeks after the third vaccine dose, we observed waning of both type-specific and cross-reactive antibodies (**Extended Data Fig. 2**). The low fractions of BA.1-specific antibodies in heterologously boosted mice at this memory phase suggest that imprinting effects persist over time after booster vaccinations. As compared to the one prime/one boost regimen, serum antibody response in mice receiving an extra priming dose encoding the Wuhan-1 spike showed greater effects of imprinting, suggesting that the number of antecedent antigen exposures shapes the serum antibody response.

As repetitive SARS-CoV-2 spike antigen exposures that occur during breakthrough infections can overcome imprinting and enhance variant-specific responses^27^, we tested whether this also happens in the context of mRNA vaccination. We immunized the two primes/one boost mice with a fourth vaccine dose of BA.1-matched vaccine (1273/1273/529/529) ten months after the third dose and collected sera 3 weeks later (**Fig. 2i**). Mice immunized with homologous mRNA-1273.529 (529/529/529/529) vaccines were included for comparison. BA.1-specific IgG titers in 1273/1273/529/529- and 529/529/529/529-immunized mice were comparable (**Fig. 2j**, *right panel*), although there was a trend towards a lower fraction of type-specific antibodies in the heterologously boosted mice (5% versus 20%, p = 0.10; **Fig. 2k**, *right panel*). Whereas all heterologously boosted mice lacked type-specific neutralizing antibodies against BA.1 after the first BA.1-matched vaccine dose (**Fig. 2h**, *right panel*), serum from 8 of 12 mice neutralized BA.1 infections with BA.1-specific antibodies after the second matched vaccine dose (**Fig. 2l**, *right panel*). Thus, in mice boosted with two doses of Omicron-matched vaccines, imprinting by antecedent vaccines can be mitigated such that variant-specific antibodies now contribute to the neutralizing response.

### Variant-specific antibodies in humans receiving a third or fourth dose of variant-matched vaccines

We next analyzed sera from individuals participating in clinical trials of B.1.351, BA.1, and BA.5 variant-matched vaccines (NCT04405076 and NCT04927065) that had no prior SARS-CoV-2 infection history as determined by negative anti-nucleocapsid antibody tests. In one study, participants were immunized with a primary two-dose series of 100 μg of mRNA-1273 and boosted at least six months later with 50 μg of mRNA-1273 (1273/1273/1273), mRNA-1273.351 (1273/1273/351), or a bivalent mRNA-1273.211 vaccine composed of a 1:1 ratio of mRNA-1273 and mRNA-1273.351 (1273/1273/211) (**Extended Data Fig. 3a**). Although Wuhan-1 spike-specific antibodies were detected in all cohorts, none of the booster vaccines induced appreciable B.1.351 spike-specific antibody responses (**Extended Data Fig. 3b-c**). Correspondingly, neutralizing activity by Wuhan-1-specific antibodies against Wuhan-1 pseudovirus was detected in all groups, but neutralization of B.1.351 pseudovirus was not observed by B.1.351 spike-specific antibodies (**Extended Data Fig. 3d**). A greater Wuhan-1-specific response was observed in 1273/1273/211 than in 1273/1273/351 participants, likely due to the inclusion of the Wuhan-1 spike-encoding mRNA in the mRNA-1273.211 booster. Despite the limited sample size, these results suggest that the binding and neutralization of B.1.351 seen across all three vaccination groups are conferred principally by cross-reactive antibodies. Moreover, the lack of B.1.351-specific antibodies in heterologously boosted participants suggests that responses were shaped by the primary vaccination series.

In a second study, individuals received two 100 μg and one 50 μg doses of mRNA-1273 and were boosted at least six months later with a fourth 50 μg dose of mRNA-1273.214, a bivalent vaccine composed of mRNA-1273 and mRNA-1273.529 (1273/1273/1273/214) (**Fig. 3a**). Sera were collected immediately before and one month after the fourth dose. Although binding reactivities against Wuhan-1 and BA.1 spike proteins were boosted by the bivalent vaccine by 2.8- and 3.1-fold, respectively (p = 0.0020 and p = 0.0039; **Fig. 3b**), we detected Wuhan-1 spike-specific and Wuhan-1/BA.1 cross-reactive but not BA.1 spike-specific antibodies (**Fig. 3c**). The fraction of Wuhan-1 spike-specific antibodies decreased slightly (24 to 19%, p = 0.023) after boosting with the bivalent vaccine, likely due to an expansion of cross-reactive antibodies (**Fig. 3d**). Functional analysis of pre- and post-boost sera revealed increases in neutralizing activity against Wuhan-1 and BA.1 pseudoviruses by 2.8- and 1.8-fold, respectively (p = 0.0020; **Fig. 3e**). Whereas inhibition of Wuhan-1 pseudovirus infection was conferred by both Wuhan-1 spike-specific and cross-reactive antibodies, neutralization of BA.1 pseudovirus was conferred exclusively by cross-reactive antibodies (**Fig. 3f**). Thus, in humans, a 50 μg of Wuhan-1/BA.1 bivalent mRNA booster vaccine following three doses of mRNA-1273 induced cross-reactive but not BA.1 spike-specific antibodies.

**Figure 3.**
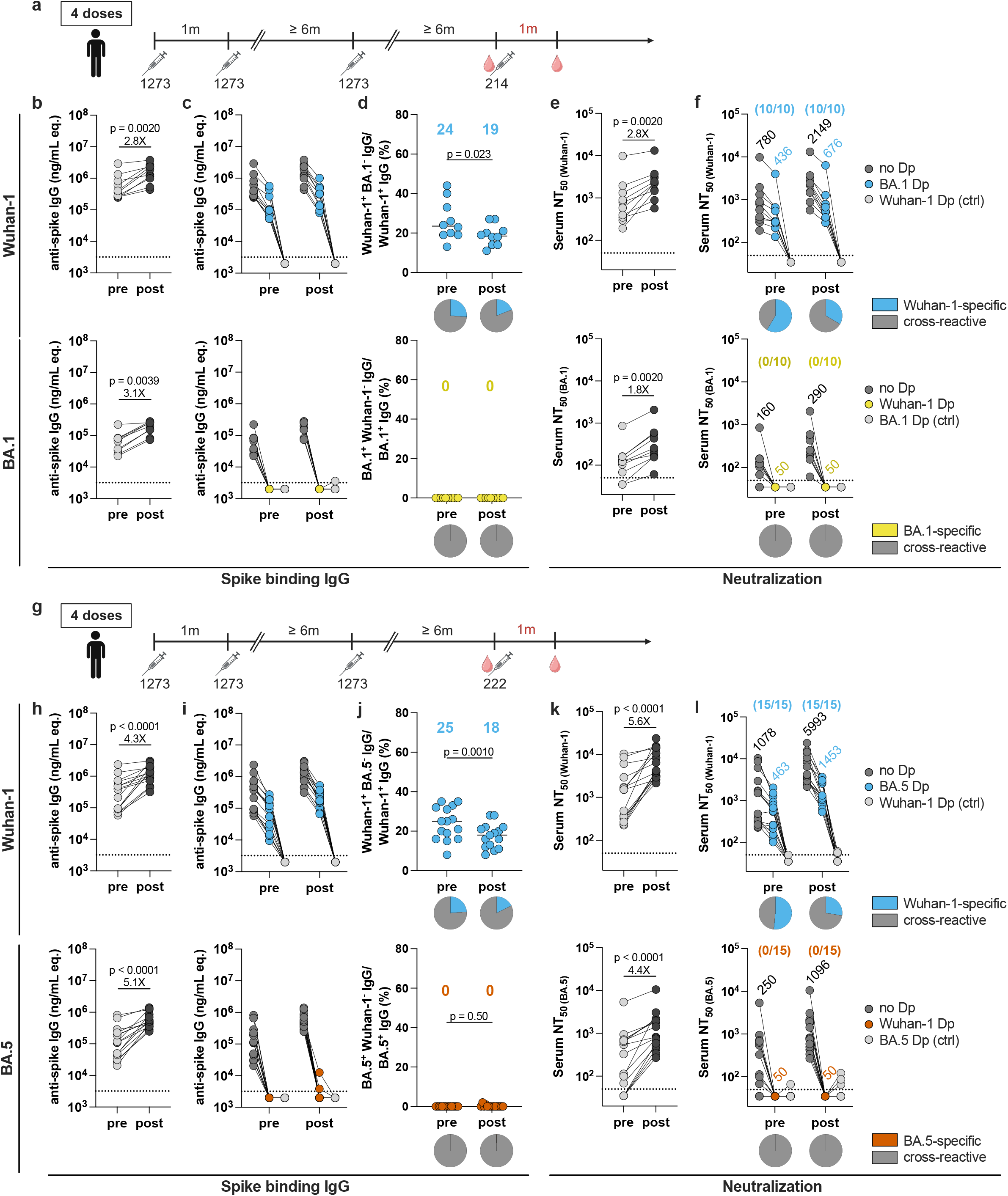
Low levels of Omicron-specific antibodies in humans boosted with a fourth dose of BA.1- or BA.5-matched vaccines. **a, g**, Schemes of immunizations and blood draws. Clinical trial participants received a primary two-dose series of 100 μg of mRNA-1273, a third dose of 50 μg of mRNA-1273, and then were boosted at least six months later with 50 μg of Wuhan-1/BA.1 bivalent vaccine (mRNA-1273.214) (**a**, n = 10) or Wuhan-1/BA.5 bivalent vaccine (mRNA-1273.222) (**g**, n = 15). Sera were collected immediately before and one month after the fourth dose. Subjects with no prior SARS-CoV-2 infection history were selected for analysis. 1273, mRNA-1273; 214, mRNA-1273.214; 222, mRNA-1273.222. **b, h**, Paired analysis of pre- and post-boost serum antibody reactivity against Wuhan-1 and BA.1 (**b**) or BA.5 (**h**) spike proteins. **c, i**, Antibody binding to Wuhan-1, BA.1 (**c**), or BA.5 (**i**) spike proteins of pre- and post- boost sera pre-cleared with empty beads (no Dp), BA.1 spike-loaded beads (BA.1 Dp), BA.5 spike-loaded beads (BA.5 Dp), or Wuhan-1 spike-loaded beads (Wuhan-1 Dp) (connecting lines represent sera from the same individual; dotted lines show the LOD; values at the LOD are plotted slightly below the LOD for visualization). **d, j**, Percentages of Wuhan-1 (*top*), BA.1 (**d**, *bottom*), and BA.5 (**j**, *bottom*) spike-specific antibodies before and after the fourth immunization. Percentages were calculated as described in Fig. 1h (numbers at the top illustrate median values). Pie charts illustrate the fraction of type-specific and cross-reactive antibodies in each group. **e, k**, Paired analysis of pre- and post-boost serum neutralizing activity against Wuhan-1, BA.1 (**e**), and BA.5 (**k**) pseudoviruses. **f, l**, Neutralizing activity of pre-cleared sera against Wuhan-1 (*top*), BA.1 (**f**, *bottom*), and BA.5 (**l**, *bottom*) pseudoviruses (numbers above data points indicate the GMT; fractions in parentheses at the top indicate the numbers of individuals with detectable neutralization by type-specific antibodies). Pie charts illustrate the fractions of neutralization mediated by type-specific and cross-reactive antibodies in each group. Serum samples with an NT_50_ below the LOD after pre-clearing with empty beads (no DP) were not used in pie chart analyses. All statistical analysis with Wilcoxon signed-rank test.

We also evaluated a separate human clinical trial group that was boosted with a fourth 50 μg dose of mRNA-1273.222, a Wuhan-1/BA.5 bivalent vaccine (1273/1273/1273/222) (**Fig. 3g**). Sera were collected immediately before and one month after the fourth dose. This bivalent vaccine increased binding titers against Wuhan-1 and BA.5 spike proteins by 4.3- and 5.1-fold, respectively (p < 0.0001; **Fig. 3h**). Whereas Wuhan-1 spike-specific and cross-reactive antibodies were observed before and after boosting, BA.5 spike-specific antibodies were detected in only 2 of 15 individuals and at low levels after boosting (**Fig. 3i**, *bottom panel*). The fraction of Wuhan-1 spike-specific antibodies in serum decreased (25 to 18%, p = 0.0010), suggesting an expansion of cross-reactive antibodies (**Fig. 3j**). Neutralizing titers against Wuhan-1 and BA.5 pseudoviruses were increased by 5.6- and 4.4-fold, respectively, after boosting (p < 0.0001; **Fig. 3k**). Whereas Wuhan-1 pseudovirus was neutralized by both Wuhan-1 spike-specific and cross-reactive antibodies, BA.5 pseudovirus was inhibited entirely by cross-reactive antibodies (**Fig. 3l**). Overall, administration of one 50 μg dose of the Wuhan-1/BA.5 bivalent mRNA vaccine following three doses of mRNA-1273 expanded cross-reactive antibodies and induced minimal levels of BA.5-specific antibodies.

### Virus-specific antibodies in humans receiving a fifth dose of XBB.1.5-matched vaccines

Omicron subvariant XBB.1.5 arose in December of 2022 and soon became prevalent in most parts of the world^28^. A clinical trial (NCT04927065) was designed to examine the immunogenicity of two XBB.1.5-matched vaccines, a bivalent BA.5/XBB.1.5 vaccine (mRNA-1273.231) and a monovalent XBB.1.5 vaccine (mRNA-1273.815). We analyzed serum samples of individuals immunized with a primary series of two 100 μg doses of mRNA-1273, boosted with a 50 μg dose of mRNA-1273, a 50 μg dose of mRNA-1273.222 (Wuhan-1/BA.5 bivalent vaccine), and then finally a 50 μg dose of BA.5/XBB.1.5 bivalent vaccine (1273/1273/1273/222/231, **Fig. 4a**) or XBB.1.5 monovalent vaccine (1273/1273/1273/222/815, **Fig. 5a**). The primary vaccine series was given one month apart, and all boosters were given at least six months after the prior immunization. None of the subjects had a history of natural SARS-CoV-2 infection and were negative for anti-nucleocapsid antibodies. Sera were collected immediately before and one month after the fifth dose.

**Figure 4.**
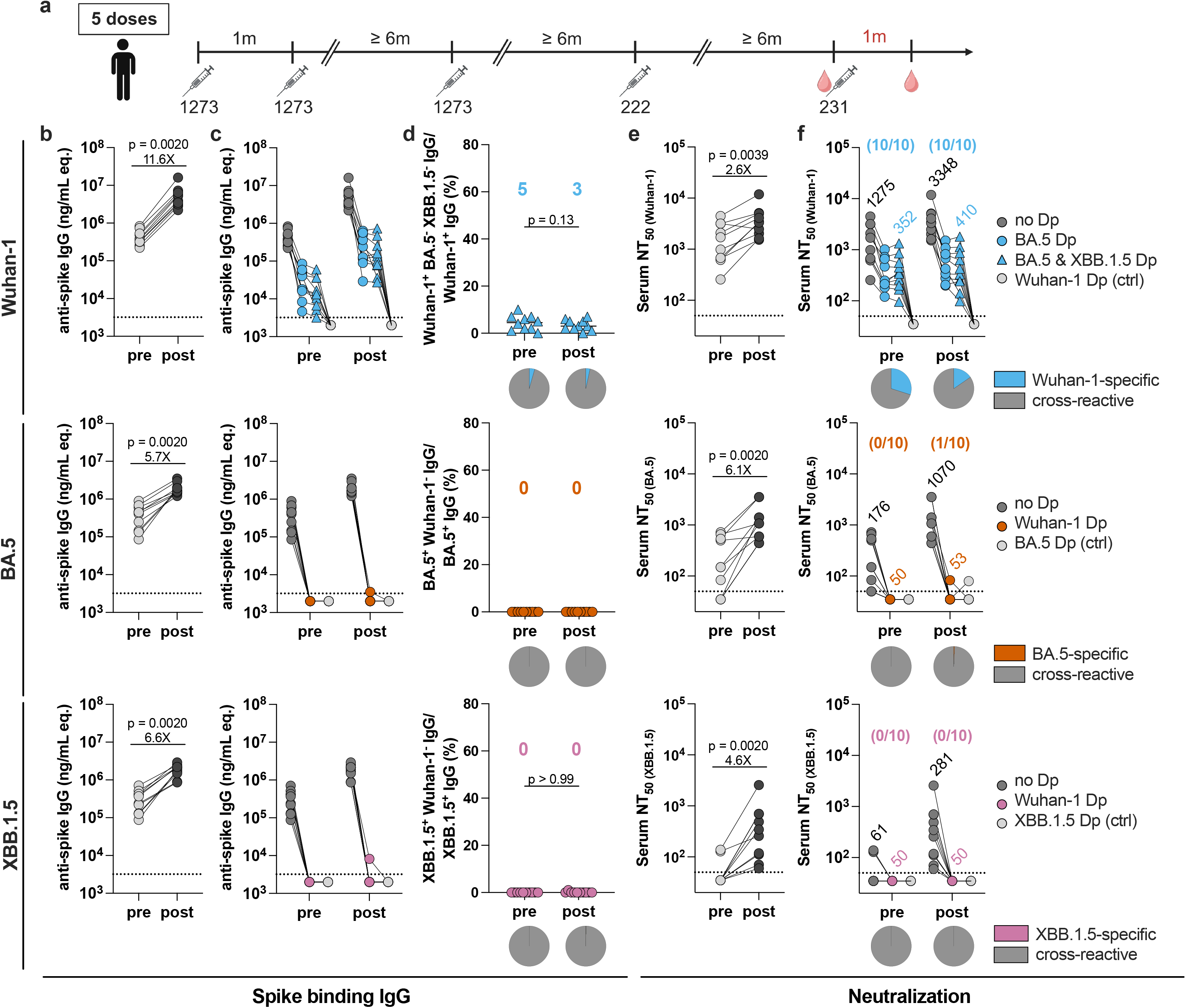
Humans administered a BA.5/XBB.1.5 bivalent vaccine after a previous BA.5-matched booster develop cross-neutralizing antibodies in serum. **a,** Scheme of immunizations and blood draws. Clinical trial participants received a primary two-dose mRNA-1273 series, a third dose of mRNA-1273, a fourth dose of mRNA-1273.222 (Wuhan-1/BA.5 bivalent vaccine), and a fifth dose of mRNA-1273.231 (BA.5/XBB.1.5 bivalent vaccine) (n = 10). Primary series vaccines were given at 100 μg doses one month apart; booster vaccines were given at 50 μg doses at least six months from the prior immunization. Sera were collected immediately before and one month after the fifth dose. Subjects with no prior SARS-CoV-2 infection history were selected for analysis. 1273, mRNA-1273; 222, mRNA-1273.222; 231, mRNA-1273.231. **b**, Paired analysis of pre- and post-boost serum antibody reactivity against Wuhan-1 (*top*), BA.5 (*middle*), and XBB.1.5 (*bottom*) spike proteins. **c**, Antibody binding to Wuhan-1, BA.5, and XBB.1.5 spike proteins of pre-cleared pre- and post-boost sera. No Dp, depletion with empty beads; BA.5 Dp, depletion with BA.5 spike-loaded beads; BA.5 & XBB.1.5 Dp, depletion with a 1:1 mixture of BA.5-loaded and XBB.1.5-loaded beads; Wuhan-1 Dp, depletion with Wuhan-1 spike-loaded beads (connecting lines represent sera from the same individual; dotted lines show the LOD; values at the LOD are plotted slightly below the LOD). **d**, Percentages of Wuhan-1-specific (BA.5- and XBB.1.5-non-reactive), BA.5-specific (Wuhan-1-non-reactive), and XBB.1.5-specific (Wuhan-1-non-reactive) antibodies before and after the fifth immunization. Percentages were calculated as described in Fig. 1h (numbers at the top illustrate median values). Pie charts illustrate the fractions of type-specific and cross-reactive antibodies. **e**, Paired analysis of pre- and post-boost serum neutralizing activity against Wuhan-1, BA.5, and XBB.1.5 pseudoviruses. **f**, Neutralizing activity of pre-cleared sera against Wuhan-1, BA.5, and XBB.1.5 pseudoviruses (numbers above data points indicate the GMT; fractions in parentheses at the top indicate the numbers of individuals with detectable neutralization by type-specific antibodies). Pie charts illustrate the fractions of neutralization mediated by type-specific and cross-reactive antibodies. Serum samples with an NT_50_ below the LOD after pre-clearing with empty beads (no DP) were not used in pie chart analyses. All statistical analysis with Wilcoxon signed-rank test.

**Figure 5.**
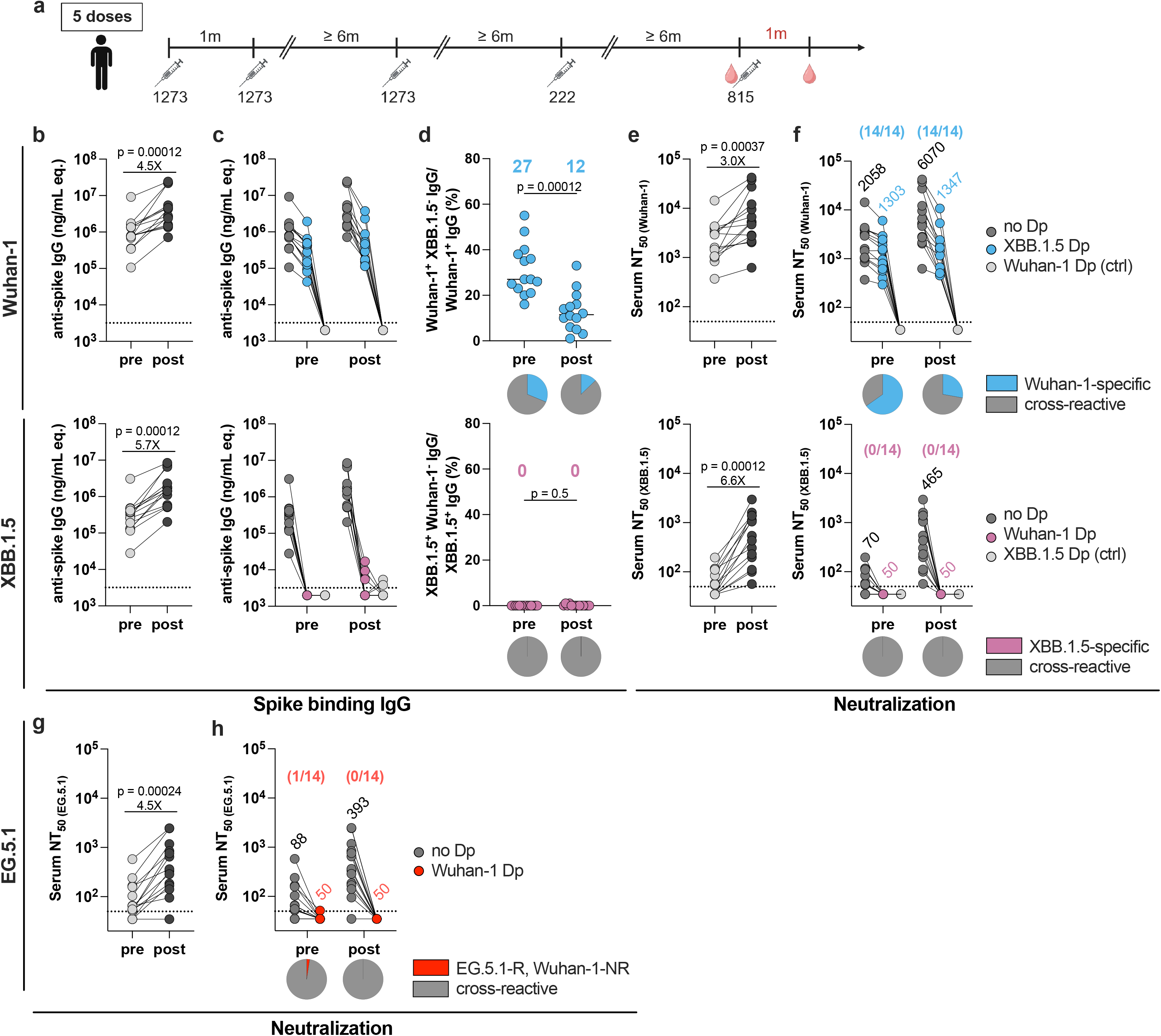
Humans administered an XBB.1.5 monovalent vaccine after a previous BA.5-matched booster develop cross-neutralizing antibodies in serum. **a**, Scheme of immunizations and blood draws. Clinical trial participants received a primary two-dose mRNA-1273 series, a third dose of mRNA-1273, a fourth dose of mRNA-1273.222 (Wuhan-1/BA.5 bivalent vaccine), and a fifth dose of mRNA-1273.815 (XBB.1.5 monovalent vaccine) (n = 14). Primary series vaccines were given at 100 μg doses one month apart; booster vaccines were given at 50 μg doses at least six months from the prior immunization. Sera were collected immediately before and one month after the fifth dose. Subjects with no prior SARS-CoV-2 infection history were selected for analysis. 1273, mRNA-1273; 222, mRNA-1273.222; 815, mRNA-1273.815. **b**, Paired analysis of pre- and post-boost serum antibody reactivity against Wuhan-1 and XBB.1.5 spike proteins. **c**, Antibody binding to Wuhan-1 and XBB.1.5 spike proteins of pre- and post-boost sera pre-cleared with empty beads (no Dp), XBB.1.5 spike-loaded beads (XBB.1.5 Dp), or Wuhan-1 spike-loaded beads (Wuhan-1 Dp) (connecting lines represent sera from the same individual; dotted lines show the LOD; values at the LOD are plotted slightly below the LOD). **d**, Percentages of Wuhan-1 and XBB.1.5 spike-specific antibodies before and after the fifth immunization. Percentages were calculated as described in Fig. 1h (numbers at the top illustrate median values). Pie charts illustrate the fractions of type-specific and cross-reactive antibodies. **e**, Paired analysis of pre- and post-boost serum neutralizing activity against Wuhan-1 and XBB.1.5 pseudoviruses. **f**, Neutralizing activity of pre- cleared sera against Wuhan-1 and XBB.1.5 pseudoviruses (numbers above data points indicate the GMT; fractions in parentheses at the top indicate the numbers of individuals with detectable neutralization by type-specific antibodies). Pie charts illustrate the fractions of neutralization mediated by type-specific and cross-reactive antibodies. Serum samples with an NT_50_ below the LOD after pre-clearing with empty beads (no DP) were not used in pie chart analyses. **g**, Paired analysis of pre- and post-boost serum neutralizing activity against EG.5.1 pseudovirus. **h**, Neutralizing activity of sera pre-cleared with Wuhan-1 spike protein against EG.5.1 pseudovirus (numbers above data points indicate the GMT; fractions in parentheses at the top indicate the numbers of individuals with detectable neutralization by type-specific antibodies). Pie charts illustrate the fractions of neutralization mediated by type-specific and cross-reactive antibodies. Serum samples with an NT_50_ below the LOD after pre-clearing with empty beads (no DP) were not used in pie chart analyses. R, reactive; NR, non-reactive. All statistical analysis with Wilcoxon signed-rank test.

Binding reactivity against Wuhan-1, BA.5, and XBB.1.5 spike proteins was boosted by the bivalent BA.5/XBB.1.5 vaccine (11.6-, 5.7-, and 6.6-fold, respectively, p = 0.0020, **Fig. 4b**). Over ∼90% of Wuhan-1 spike-reactive IgG before and after the fifth dose cross-reacted with the BA.5 spike protein, as pre-clearing with BA.5 spike-coated beads decreased Wuhan-1 spike reactivity by approximately 10-fold (**Fig. 4c**). To determine whether the residual Wuhan-1 spike-reactive antibodies were Wuhan-1-specific or cross-reactive to XBB.1.5 spike, we co-depleted sera with BA.5 and XBB.1.5 spike proteins (**Fig. 4c-d**). The remaining Wuhan-1 spike-reactive antibodies were type-specific and did not cross-react with BA.5 or XBB.1.5. Despite the increased serum antibody reactivity to BA.5 and XBB.1.5 spike proteins after the BA.5/XBB.1.5 bivalent booster, BA.5 or XBB.1.5 spike-specific antibodies were not appreciably induced. The BA.5/XBB.1.5 bivalent vaccine also boosted neutralizing responses against Wuhan-1, BA.5, and XBB.1.5 pseudoviruses (2.6-, 6.1-, and 4.6-fold, respectively; p = 0.0039, 0.0020, and 0.0020, **Fig. 4e**). Depletion analysis revealed that over 70% of serum neutralizing activity against Wuhan-1 was mediated by cross-reactive antibodies (**Fig. 4f**), whereas the inhibition of BA.5 and XBB.1.5 infections was conferred entirely by antibodies that cross-reacted with Wuhan-1 spike. These data suggest an imprinting effect of the antecedent mRNA vaccines, in which two exposures to Omicron subvariant spike proteins after three exposures to the Wuhan-1 spike resulted in expansion of cross-reactive, cross-neutralizing antibodies that target the XBB.1.5 variant. In humans, this regimen was insufficient to elicit appreciable levels of Omicron-specific antibodies.

We next evaluated serum from individuals that received a fifth dose of mRNA-1273.815, the currently administered monovalent mRNA vaccine booster targeting the XBB.1.5 spike. Antibody titers against Wuhan-1 and XBB.1.5 spike proteins were increased by 4.5- and 5.7-fold, respectively (p = 0.00012, **Fig. 5b**). The fraction of Wuhan-1 spike-specific antibodies decreased after the booster dose (27 to 12%, p = 0.00012), again consistent with an expansion of cross-reactive antibodies (**Fig. 5c-d**). XBB.1.5 spike-specific antibodies were absent before mRNA-1273.815 boosting and detected in only 3 of 14 individuals at low levels after boosting, indicating that the increased reactivity to XBB.1.5 spike was principally mediated by cross-reactive antibodies. The XBB.1.5 monovalent booster also enhanced neutralizing activity against Wuhan-1 and XBB.1.5 pseudoviruses, by 3.0- and 6.6-fold, respectively (p = 0.00037 and p = 0.00012, **Fig. 5e**). Whereas a fraction of Wuhan-1 neutralization before and after boosting was conferred by Wuhan-1-specific antibodies, neutralizing activity against XBB.1.5 was mediated entirely by antibodies that cross-reacted with Wuhan-1 spike (**Fig. 5f**).

In August of 2023, XBB.1.5 infections were superseded by subvariants of the EG.5.1 lineage^28^. To test whether responses to the monovalent XBB.1.5 vaccine had neutralizing activity against EG.5.1, we incubated sera from 1273/1273/1273/222/815-vaccinated subjects with EG.5.1 pseudovirus. Inhibitory activity against EG.5.1 infection was boosted by 4.5-fold (p = 0.00024, **Fig. 5g**), and neutralization was abolished in sera pre-cleared with Wuhan-1 spike, suggesting that inhibitory activity against EG.5.1 was mediated by Wuhan-1 cross-reactive antibodies (**Fig. 5h**). Overall, these data from human clinical cohorts suggest that monovalent XBB.1.5-matched mRNA vaccine given as a fifth dose boosted cross-reactive neutralizing antibodies that were shaped by the original mRNA-1273 vaccine.

### Breadth and specificity of cross-reactive serum neutralizing antibody responses after boosting

To begin to define the epitopes recognized by cross-neutralizing antibodies from mRNA-1273.815-boosted individuals (**Fig. 5a**), we developed a bead-based mAb competition assay (**Extended Data Fig. 4a**). XBB.1.5 spike-coated beads were incubated first with serum samples and then, without washing, fluorescently tagged neutralizing mAbs that bind to mapped epitopes were added for detection by flow cytometry. We examined binding of 8 published and unpublished XBB.1.5 cross-reactive neutralizing mAbs that recognize previously defined RBD antigenic sites^29^ (class 1: rCOV2-3731; class 3: S309, rCOV2-3872, rCOV2-3967, and rCOV2-3889; class 4: rCOV2-4030) and the S2 stem helix region (CC40.8 and CV3-25) ^19,25,30^. Competition was evaluated by measuring reductions in the mean fluorescence intensity associated with mAb binding. Naïve sera from individuals with no prior SARS-CoV-2 infection or vaccination history were included as negative controls. Serum cross-reactive antibodies from most 1273/1273/1273/222/815-vaccinated individuals bound at or near all tested neutralizing epitopes in XBB.1.5 RBD and S2 and were boosted to similar extents (**Extended Data Fig. 4b-c**). This analysis suggests that the XBB.1.5 monovalent vaccine booster induced antibodies that recognize multiple cross-reactive and neutralizing epitopes on the spike protein.

Given that some neutralizing epitopes on the SARS-CoV-2 spike protein are evolutionarily conserved^31,32^, we hypothesized that antibodies from the serum of XBB.1.5-boosted individuals might neutralize more distantly related coronaviruses. Accordingly, we tested the sera for inhibitory activity to pseudoviruses encoding the spike protein of pangolin coronavirus Guangdong (Pang/GD, 67.9, 96.4, 97.6, and 89.8% sequence identity in NTD, RBD, S2, and spike compared to SARS-CoV-2 Wuhan-1), Severe acute respiratory syndrome coronavirus 1 (SARS-CoV-1, 47.4, 73.1, 88.4, and 76.0% sequence identity in NTD, RBD, S2, and spike compared to SARS-CoV-2 Wuhan-1), and Middle East respiratory syndrome coronavirus (MERS-CoV, 16.3, 18.1, 38.0, and 28.6% sequence identity in NTD, RBD, S2, and spike compared to SARS-CoV-2 Wuhan-1) (**Extended Data Fig. 5**). We also modified our depletion assay to determine the contribution of cross-reactive antibodies that engaged the N-terminal domain (NTD), RBD, or S2 protein to neutralization. We first compared the effect of depletion of Wuhan-1 NTD, RBD, or S2 targeting antibodies on neutralization of Wuhan-1 or XBB.1.5 pseudoviruses (**Fig. 6a-b** and **Extended Data Fig. 6a**). Neutralization of Wuhan-1 pseudovirus was detected in serum before and after XBB.1.5-boosting, and this activity was mediated principally by anti-RBD antibodies (12.6-fold reduction in GMT after RBD depletion) compared to anti-NTD (1.8-fold reduction) or anti-S2 (no reduction) antibodies (**Fig. 6a**, *right panel*). In comparison, pre-boost serum had poor inhibitory activity against XBB.1.5. However, after boosting, serum neutralized XBB.1.5 pseudovirus infection, and this activity was mediated exclusively by anti-RBD antibodies (**Fig. 6b**, *right panel*).

**Figure 6.**
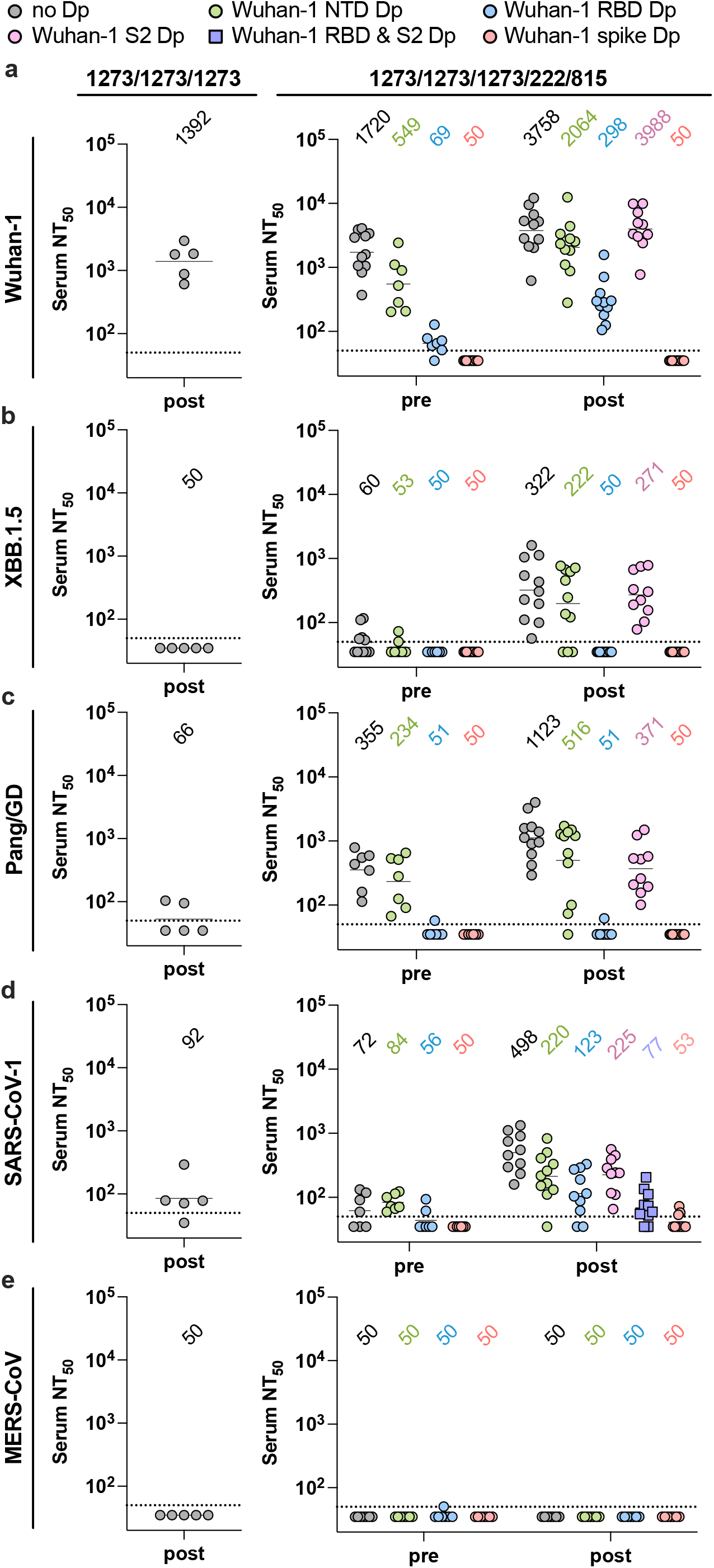
Neutralizing activity of vaccine-derived serum against distantly related sarbecoviruses. 1273/1273/1273 post-boost sera (*left panels*, also described in **Extended Data Fig. 3a**) or pre- and post-boost sera relative to immunization with mRNA-1273.815 (XBB.1.5 monovalent vaccine, *right panels*, also described in Fig. 5a) were pre-cleared with Wuhan-1 NTD, RBD, S2, RBD together with S2 (RBD & S2), or spike protein. Pre-cleared sera were tested for neutralizing activity against pseudoviruses expressing Wuhan-1 (**a**), XBB.1.5 (**b**), Pang/GD (**c**), SARS-CoV-1 (**d**), or MERS-CoV (**e**) spike proteins (numbers at the top indicate GMT values; dotted lines show the LOD; values at the LOD are plotted slightly below the LOD).

Although serum obtained before or after XBB.1.5 boosting had inhibitory activity against Pang/GD, neutralizing titers were increased 3.2-fold (p = 0.037) at one month after boosting with XBB.1.5 mRNA vaccine (**Fig. 6c**, *right panel*). In comparison, individuals who received multiple doses of only mRNA-1273 showed little to no inhibitory activity against Pang/GD one month after the last dose, suggesting that heterologous boosting expands the pool of broadly cross-neutralizing antibodies (**Fig. 6c**, *left panel*). Like results with XBB.1.5, neutralization of Pang/GD was mediated almost entirely by Wuhan-1 RBD-targeting antibodies in most individuals. Serum obtained before XBB.1.5 boosting or 1 month after multiple mRNA-1273 immunizations poorly neutralized SARS-CoV-1, a more distantly related sarbecovirus (**Fig. 6d**). One month after XBB.1.5 mRNA vaccine boosting, neutralization of SARS-CoV-1 was observed, and this was mediated by spike-reactive antibodies imprinted by the mRNA-1273 vaccines. Pre-clearing of sera with Wuhan-1 RBD only partially reduced SARS-CoV-1 neutralization, suggesting a less dominant role of RBD-targeting antibodies in the protection against this more distantly related coronavirus. However, co-depletion of serum antibodies with both Wuhan-1 RBD and S2 proteins reduced SARS-CoV-1 neutralization almost to baseline, which indicates significant contributions of both anti-RBD and anti-S2 antibodies to neutralization (**Extended Data Fig. 6b**). Although boosting with XBB.1.5 enhanced neutralization against two other sarbecoviruses, inhibitory activity was not detected against the distantly related merbecovirus, MERS-CoV (**Fig. 6e**). Collectively, our results suggest that the XBB.1.5 monovalent vaccine booster enabled or increased neutralizing activity against multiple sarbecoviruses, and this inhibitory activity is imprinted by antibodies targeting the Wuhan-1 RBD and S2 regions. While imprinted anti-RBD antibodies have a major role in neutralizing SARS-CoV-2 and the antigenically closer Pang/GD, imprinted anti-S2 antibodies contribute more to serum neutralizing titers against the more antigenically distant SARS-CoV-1.

## DISCUSSION

We used antibody depletion and multiplexed spike-binding detection assays to distinguish type-specific and cross-reactive serum antibody responses against SARS-CoV-2 spike proteins. Rather than performing repertoire analysis of monoclonal antibodies derived from vaccine-induced MBCs to assess the effects of imprinting, we profiled the type-specificity and cross-reactivity of serum antibodies, which have been shown to correlate with protection against infection and disease^33^. In mice, the induction of BA.1-specific neutralizing antibodies by BA.1- matched boosters was present but limited by imprinting, and this effect varied depending on the number of antigen exposures with the original and the boosting antigen. In humans without a history of SARS-CoV-2 infection, we detected little to no variant-specific serum antibody response in clinical trial participants primed with two or three doses of mRNA-1273 and boosted with B.1.351-, BA.1-, BA.5-, or XBB.1.5-matched monovalent or bivalent vaccines, suggesting strong imprinting effects of antecedent mRNA vaccinations. Serum antibody responses in humans that received Omicron-matched boosters were characterized by cross-reactive spike binding and cross-neutralizing activity that extended to distantly related members of the sarbecovirus subgenus and were likely imprinted by B cells that recognized Wuhan-1 spike.

Imprinting after SARS-CoV-2 infection and vaccination has been suggested by studies that analyze the differential reactivity of serum and MBCs to different spike proteins^2,9–11,13–17^. Although a standard analysis of serum can demonstrate whether spike-binding and neutralizing antibodies against a given variant are induced after infection or boosting, these studies do not address the composition and functional contribution of type-specific versus cross-reactive antibodies. Our spike-based depletion assay showed little to no variant-specific antibodies in serum of clinical trial participants boosted with variant-matched vaccines after priming with mRNA-1273, even when two successive doses of variant vaccines were administered. Our data showing that the spike binding and neutralizing response against Omicron strains is almost entirely cross-reactive with the Wuhan-1 spike suggest that in humans, serum antibody responses are imprinted by antecedent mRNA vaccinations. Consistent with these results, a prior study that used antibody depletion with RBD proteins reported undetectable levels of BA.5 RBD-specific antibodies in the serum of individuals boosted with Wuhan-1/BA.5 bivalent mRNA vaccines^34^. Our results are relevant to a recent study, which suggested that repeated Omicron infections could override ancestral SARS-CoV-2 immune imprinting in the serum and B cell compartments^27^. In that publication, individuals received two doses of the inactivated whole virus CoronaVac vaccine and then experienced two Omicron breakthrough infections. Differences in the immune stimulus of the initial vaccine (CoronaVac versus mRNA-1273) or subsequent Omicron exposures (breakthrough infections versus mRNA vaccines) could dictate whether cross-reactive memory B cells or *de novo* type-specific naïve B cells are activated to produce antibodies. Consistent with our findings, a preprint described imprinting of human plasma antibody responses^35^. Individuals who had received multiple vaccine doses with or without known infection history and then were boosted with the XBB.1.5 mRNA monovalent vaccine had neutralizing antibodies against current variants that were dominated by recall of pre-existing memory B cells induced by the Wuhan-1 spike. One caveat of the preprint study is that plasma samples were obtained at relatively early time points (only 7 to 13 days) after XBB.1.5 vaccine boosting, which may have preceded secondary germinal center reactions.

The imprinting effect we observed also agrees with analysis of MBCs in non-human primates, where boosting with the BA.1-matched mRNA vaccine expanded the Wuhan-1/BA.1 cross-reactive MBC population, but not BA.1 spike-specific MBCs^11^. B cell repertoire analysis in humans also showed few (< 1%) BA.1-specific MBCs in peripheral blood of individuals boosted with the BA.1-matched monovalent mRNA vaccine after priming with multiple doses of mRNA vaccines encoding Wuhan-1 spike protein^16^. While the MBC repertoire does not always reflect the serum antibody composition due to possible discrepancies in the antigen specificity between MBCs and LLPCs^18^, in the context of SARS-CoV-2 mRNA vaccine boosting, substantial imprinting effects likely occur in both memory cell compartments. Although secondary germinal centers are formed after boosting and include newly recruited naïve B cells^36^, for most individuals who had multiple prior mRNA vaccine exposures, antibodies derived from these *de novo* activated B cells contributed to a minority of the serum response against Omicron variants, at least at the time points we examined. This result might differ if the initial antigen exposure is immunologically weaker, or the subsequent exposure (*i.e*., breakthrough infection) is substantially stronger.

In mice, type-specific serum antibodies were elicited by variant-matched mRNA booster vaccines, although the degree of immune imprinting was determined in part by the number of primary and subsequent homologous and heterologous antigen exposures. Indeed, a recent fate mapping study in mice primed once with Wuhan-1 spike mRNA vaccine and heterologously boosted twice with BA.1 spike mRNA vaccine identified *de novo* serum antibodies derived from newly recruited B cells that had not participated in the original germinal center reaction^37^. Our experiments in mice also show that while repeated administration of the prototype mRNA-1273 vaccine promotes imprinting, administration of multiple boosters encoding variant spike proteins can mitigate imprinting and promote responses that yield type-specific neutralizing antibodies in serum. These findings agree with results observed with repeated influenza booster vaccines^38^. It remains to be determined how many doses of variant-matched vaccines (or breakthrough infections) are required to overcome mRNA-1273 vaccine-induced imprinting to elicit type-specific antibody responses in humans that contribute to the serum neutralizing activity. This outcome likely depends on multiple variables including the priming and boosting intervals, the number of doses of prototype mRNA-1273 vaccine administered (as suggested by our findings in one prime/one boost versus two primes/one boost mouse cohorts), and the antigenic distance of the priming and boosting antigens. Indeed, increasing the dose of antigen in booster doses enhanced antibody titers against specific influenza antigens^39^, although it remains unclear if this was due to greater induction of variant type-specific or cross-reactive antibodies.

Despite the limited induction of variant-specific serum antibodies by humans immunized and boosted with mRNA vaccines, our studies show that the cross-reactive antibodies expanded by boosting exhibited broad spike-reactivity and mediated neutralization of the priming strain, the boosting strain, subsequent circulating Omicron variants (*e.g*., EG.5.1), and even distantly related sarbecoviruses including Pang/GD and SARS-CoV-1. In this context, immune imprinting by SARS-CoV-2 mRNA vaccines does not constitute original antigenic “sin”, as cross-reactive antibodies are likely beneficial for immunity and protection. Although the sequence identity of spike protein between Wuhan-1 and XBB.1.5 strains (96.7%) versus Wuhan-1 and Pang/GD (89.8%) or SARS-CoV-1 (76.0%) is greater, some individuals had detectable yet low levels of neutralizing antibody against Pang/GD and SARS-CoV-1 but little or none against XBB.1.5 and EG.5.1 prior to XBB.1.5 mRNA vaccine boosting. Thus, despite the greater spike sequence identity between Wuhan-1 and Omicron lineage variants than Pang/GD and SARS-CoV-1, the antigenic distance is greater. This result is likely because Omicron strains have evolved multiple key substitutions in RBD and S2 epitopes that evade neutralizing antibody responses induced by Wuhan-1 spike directed vaccines.

Using competition binding studies with broadly neutralizing antibodies that recognize distinct sites in the RBD (class 1, 3, and 4 epitopes) or S2 proteins, our results suggest that boosting with XBB.1.5 mRNA vaccine expands cross-reactive antibodies that bind multiple neutralizing epitopes. However, these experiments cannot define the precise footprint of antibody recognition, as competition could occur through binding of epitopes that are proximal to neutralizing sites. More definitive epitope identification of cross-neutralizing epitopes could be informed by studies using landscape mutagenesis in the context of pseudoviruses^40,41^ or polyclonal antibody epitope mapping and cryo-electron microscopy^42,43^. Our depletion studies with Wuhan-1 NTD, RBD, and S2 suggest that the class of cross-neutralizing antibody may differ depending on the individual sarbecovirus. Whereas neutralizing responses against XBB.1.5 and Pang/GD were largely depleted by RBD, those against SARS-CoV-1 required pre-clearing with both RBD and S2 proteins to reduce inhibitory responses.

We note several limitations in our study. (a) Our study did not address serum antibody responses in the context of hybrid immunity; (b) We characterized the serum antibody responses following immunization with only mRNA vaccines against SARS-CoV-2; (c) Our studies were limited by the interval and dosing regimen of booster vaccinations used; and (d) We examined the post-boost response at a single 1-month time point in most cohorts. Analysis of serum at later time points could reveal whether variant-specific responses emerge after sustained germinal center reactions^44^.

In summary, our study shows that in humans, variant-matched mRNA boosters administered after prototype mRNA-1273 vaccinations primarily expand Wuhan-1-cross-reactive antibodies. Minimal variant-specific serum antibody responses were detected in individuals boosted with Omicron-matched mRNA vaccines, due to imprinting effects of two or more doses of the prototype vaccine. Despite the strongly imprinted serum responses by prior mRNA-1273 immunization, heterologous mRNA vaccine boosters induced cross-reactive antibodies with broadly neutralizing activity that likely contribute to their continued efficacy against severe SARS-CoV-2 disease. Intriguingly, administration of future boosters that continue to drift from the original Wuhan-1 spike might result in the further expansion of broadly neutralizing anti-RBD and S2 antibodies against multiple and even more distantly related sarbecoviruses.

## Acknowledgements

This study was supported by the National Institutes of Health (R01 AI157155, NIAID Centers of Excellence for Influenza Research and Response (CEIRR) contracts 75N93021C00014 and 75N93019C00051, to M.S.D.) and a sponsored Research Agreement with Moderna. We thank S. Mackin and B. Ying for assistance with some of the animal studies, and L. Lu for assistance with the production of some of the chimeric VSV pseudoviruses. Some of the Figure graphics were created using BioRender.

## Author contributions

C.Y.L. and S.R. established the multiplexed spike-binding assay and antibody depletion assay. S.R., S.J.Z. and J.E.C. produced the human mAbs. C.Y.L. performed mouse experiments, depletion experiments, multiplexed binding experiments, ELISA experiments, and mAb competition experiments. C.Y.L. established and performed pseudovirus neutralization assays. Z.L., Y.L., and S.P.J.W. designed and generated the VSV pseudoviruses. J.B.C. performed spike sequence alignment analysis. S.M.E. provided mRNA vaccines. G.A.A. provided SARS-CoV-2 NTD, RBD, and spike proteins. C.M.A. and J.S.M. provided Wuhan-1 S2 protein. Moderna, Inc. provided human clinical trial sera. M.S.D. supervised the research. C.Y.L. and M.S.D designed studies and wrote the initial draft, with all other authors providing editorial comments.

## EXTENDED DATA FIGURE LEGENDS

**Extended Data Figure 1. Gating strategy and validation of the multiplexed detection and antibody depletion assays. a**, Gating for the multiplexed spike-binding assay. After gating on singlets, beads pre-bound with the spike protein of different SARS-CoV-2 strains were separated based on their intensity of SPHERO™ yellow fluorophore. **b**, Pearson’s correlation between the spike-binding and ELISA endpoint titers of the mAbs evaluated in **Fig. 1c-d**. R^2^ and p values are indicated. **c**, Antibody binding of mAb mixtures pre-cleared with empty depletion beads (no Dp), B.1.351 spike-loaded beads (B.1.351 Dp), or Wuhan-1 spike-loaded beads (Wuhan-1 Dp) to Wuhan-1 spike-coated detection beads (values obtained after interpolating with CV3-25 standard curves are shown in Fig. 1f).

**Extended Data Figure 2. Persistent effects of immune imprinting in the serum of mice receiving heterologous mRNA vaccines. a**, Scheme of immunization regimen and blood draw. Mice were immunized one month apart and boosted with a third dose of mRNA vaccine three months later (n = 10, two experiments, also in Fig. 2e). Sera were collected 15 weeks after the last dose. 1273, mRNA-1273; 529, mRNA-1273.529. **b**, Antibody binding of sera pre-cleared with empty beads (no Dp), BA.1 spike-loaded beads (BA.1 Dp), or Wuhan-1 spike-loaded beads (Wuhan-1 Dp) to Wuhan-1 (*left*) and BA.1 (*right*) spike proteins (connecting lines represent sera from the same mouse; dotted lines show the LOD; values at the LOD are plotted slightly below the LOD). **c**, Percentages of Wuhan-1 (*left*) and BA.1 (*right*) spike-specific antibodies. Percentages were calculated as described in Fig. 1h (numbers at the top illustrate median values). Pie charts illustrate the fractions of type-specific and cross-reactive IgG in each group. **d, e**, Paired analysis of 3- and 15-week post-boost undepleted sera (**d**, 15-week data also shown in **b** “no Dp”) and type-specific (**e**, 15w data also shown in **b** “BA.1 Dp” or “Wuhan-1 Dp”) antibody reactivity against Wuhan-1 (*left*) and BA.1 (*right*) spike proteins (numbers on the top indicate GMT fold reduction). **c**, Kruskal-Wallis ANOVA with Dunn’s post-test.

**Extended Data Figure 3. Absence of B.1.351 spike-specific antibodies in humans receiving B.1.351-matched booster as a third-dose vaccine. a**, Scheme of immunizations and blood draw. Participants received two doses of 100 μg of mRNA-1273 one month apart and were boosted with 50 μg of the indicated mRNA vaccine at least six months later. Sera were collected one month after the third dose (n = 5). 1273, mRNA-1273; 351, mRNA-1273.351; 211, mRNA-1273.211 (Wuhan-1/B.1.351 bivalent vaccine). **b**, Antibody binding to Wuhan-1 (*top*) and B.1.351 (*bottom*) spike proteins of sera pre-cleared with empty beads (no Dp), B.1.351 spike-loaded beads (B.1.351 Dp), or Wuhan-1 spike-loaded beads (Wuhan-1 Dp) (connecting lines represent sera from the same individual; dotted lines show the LOD; values at the LOD are plotted slightly below the LOD). **c**, Percentages of Wuhan-1 (*top*) and B.1.351 (*bottom*) spike-specific antibodies. Percentages were calculated as described in Fig. 1h (numbers at the top illustrate median values). Pie charts illustrate the fractions of type-specific and cross-reactive IgG in each group. **d**, Neutralizing activity of pre-cleared sera against Wuhan-1 (*top)* and B.1.351 (*bottom*) pseudoviruses (numbers above data points indicate the GMT; fractions in parentheses at the top indicate the numbers of individuals with detectable neutralization by type-specific antibodies). Pie charts illustrate the fractions of neutralization mediated by type-specific and cross-reactive antibodies. Serum samples with an NT_50_ below the LOD after pre-clearing with empty beads (no DP) were not used in pie chart analyses. **c**, Kruskal-Wallis ANOVA with Dunn’s post-test.

**Extended Data Figure 4. XBB.1.5 monovalent booster increases serum binding to multiple neutralizing epitopes on the XBB.1.5 spike protein. a**, Schematic of the mAb competition assay (competing serum antibody, red; non-competing serum antibody, salmon; fluorescence-labeled mAb, blue). **b**, Percent binding of the indicated mAbs competed by serially diluted pre- and post-boost serum of XBB.1.5 monovalent vaccine boosted individuals described in Fig. 5 to XBB.1.5 spike protein. Two naïve sera (black curves, no SARS-CoV-2 infection or vaccination history) were included as control. **c**, Competition titer of pre- and post-boost sera that inhibits 50% binding of the indicated mAbs (numbers on the top indicate GMT fold increases), derived from **b**.

**Extended Data Figure 5. Spike protein sequence alignment.** Multiple sequence alignment of SARS-CoV-1 (P59594, UNIPROT), Pang/GD (EPI_ISL_410721, GISAID), XBB.1.5 (EPI_ISL_16713058, GISAID), and Wuhan-1 (EPI_ISL_402124, GISAID) spike proteins. The Wuhan-1 spike is shown in the last row with relative variant sequence changes indicated. The color ribbons beneath the sequence correspond to the specific regions of the spike protein (NTD, green; RBD, blue; S2, pink).

**Extended Data Figure 6. Depletion of Wuhan-1 NTD, RBD, S2, and spike-reactive antibodies from serum. a**, Antibody binding to Wuhan-1 NTD, RBD, S2, or spike proteins of pre- and post-boost sera of 1273/1273/1273/222/815-vaccinated individuals pre-cleared with empty beads (no Dp), Wuhan-1 N-terminal domain protein-loaded beads (NTD Dp), RBD-loaded beads (RBD Dp), S2 protein-loaded beads (S2 Dp), a 1:1 mixture of RBD-loaded and S2-loaded beads (RBD & S2 Dp), or spike-loaded beads (spike Dp) (dotted lines show the LOD; values at the LOD are plotted slightly below the LOD). Binding to Wuhan-1 NTD, RBD, and S2 was interpolated from SARS2-57, SARS2-38, and CV3-25 standard curves, respectively. **b**, Serum neutralizing activity of post-boost sera pre-cleared with empty beads (no Dp), Wuhan-1 RBD-loaded beads (RBD Dp), or a mixture of Wuhan-1 RBD-loaded and S2-loaded beads (RBD & S2 Dp) against SARS-CoV-1 pseudovirus (data also shown in Fig. 6d, *right*) (connecting lines represent sera from the same individual). Statistical analysis: Wilcoxon signed-rank test.

## METHODS

### Cells and viruses

Vero-TMPRSS2 cells^45^ were cultured at 37°C in Dulbecco’s Modified Eagle medium (DMEM) supplemented with 10% fetal bovine serum (FBS), 10 mM HEPES pH 7.3, 1 mM sodium pyruvate, 1X nonessential amino acids, 100 U/ml of penicillin–streptomycin, and 5 μg/mL of blasticidin. Chimeric VSV pseudoviruses expressing spike proteins corresponding to the ancestral SARS-CoV-2 Wuhan-1 virus, variant strains B.1.351, BA.1, BA.5, XBB.1.5, and EG.5.1, and other coronaviruses including Pang/GD, SARS-CoV-1, and MERS-CoV were produced according to published methods^26^. Briefly. the spike genes of different coronaviruses including SARS-CoV-2 (Wuhan-1, GenBank MN908947.3; B.1.351, GenBank OX008568.1; BA.1, GenBank OR321077.1; BA.5, GenBank QY979334.1; XBB.1.5, GenBank OP790748.1; and EG.5.1, GenBank OX463749.1), Pang/GD (GenBank MT799524.1), SARS-CoV-1 (GenBank JN854286.1), and MERS-CoV (GenBank MK357909.1) were synthesized (Integrated DNA Technologies) and inserted into an infectious molecular clone of VSV^46,47^ as described previously^26^. Modifications to the cytoplasmic tail (21 amino acid deletion) of the spike genes were generated to promote secretion. Other VSV N, P, L and G expression plasmids have been described^47^.

Recovery of recombinant VSV was performed as described^47^. BSRT7/5 cells^48^ were inoculated with vaccinia virus vTF7-3 and subsequently transfected with T7-expression plasmids encoding VSV N, P, L, and G, and an antigenomic copy of the viral genome. Cell culture supernatants were collected at 56-72 h, clarified by centrifugation at 1,000 x *g* for 5 min, and filtered through a 0.22 μm filter. Virus was plaque-purified on Vero CCL81 cells in the presence of 25 μg/mL of cytosine arabinoside (Sigma-Aldrich), and plaques in agarose plugs were amplified on Vero CCL81 cells. Viral stocks were amplified on MA104 cells at an MOI of 0.01 in Medium 199 containing 2% FBS and 20 mM HEPES pH 7.7 at 34°C. Viral supernatants were harvested upon extensive cytopathic effect and clarified of cell debris by centrifugation at 1,000 x g for 5 min. Aliquots were stored at −80°C.

### Animal experiments

Animal studies were carried out in accordance with the recommendations in the Guide for the Care and Use of Laboratory Animals of the National Institutes of Health. Protocols were approved by the Institutional Animal Care and Use Committee at the Washington University School of Medicine (assurance number A3381–01). Six-week-old female C57BL/6J mice purchased from The Jackson Laboratory (Cat. #000664) were used for all experiments. Animals were housed in groups of five and fed standard chow diets. Photoperiod was 12 h on:12 h off dark/light cycle. The ambient animal room temperature was 70°F, controlled within ± 2°F. The room humidity was 50%, controlled within ± 5%. Mice described in **Fig. 1g-h** were primed via intraperitoneal injection with 0.25 μg and boosted with 1 μg of mRNA vaccines one month apart, and serum was collected 3 weeks later. Mice described in Fig. 2 were immunized according to the depicted schemes. The primary vaccines were given 4 weeks apart, the third dose vaccine was given 3 months later, and the fourth dose vaccine was given 10 months after the third dose. All vaccines given to mice are preclinical materials and were administered intraperitoneally at 0.25 μg mRNA per mouse. Serum samples were collected 3 weeks after the final immunization.

### Clinical study design and human participants

Human samples were obtained from open-label phase 2/3 studies (NCT04405076 and NCT04927065) that evaluate the immunogenicity, safety, and reactogenicity of mRNA booster vaccines in adults who had already received 2-dose primary series (100 µg) of mRNA-1273. Participants were enrolled in a sequential, non-randomized manner and received boosters of 50 µg of mRNA-1273, mRNA- 1273.351, or mRNA-1273.211 (cohort 1, **Extended Data** Fig. 3), 50 µg of mRNA-1273 followed by 50 µg of mRNA-1273.214 or mRNA-1273.222 (cohort 2, Fig. 3), or 50 µg of mRNA-1273 followed by 50 µg of mRNA-1273.222 and then 50 µg of mRNA-1273.231 (cohort 3, Fig. 4), or 50 µg of mRNA-1273 followed by 50 µg of mRNA-1273.222 and then 50 µg of mRNA-1273.815 (cohort 4, Fig. 5).

For participants in trial NCT04927065 boosted with a fifth dose of mRNA-231 or mRNA- 1273.815, median (interquartile ranges) intervals were 8.2 (7.8-9.8) and 9.0 (7.7-11.5) months between the 2^nd^ and 3^rd^ doses (both mRNA-1273), 9.8 (8.3-10.3) and 9.2 (8.2-10.3) months between the 3^rd^ and 4^th^ doses (mRNA-1273 and mRNA-1273.222, respectively), and 8.3 (8.1- 8.4) and 8.2 (8.1-8.3) months between the 4^th^ and 5^th^ doses, respectively. Interim 29-day analysis results are reported.

The trials are being conducted across 23 sites in the United States of America, in accordance with the International Council for Harmonisation of Technical Requirements for Registration of Pharmaceuticals for Human Use, Good Clinical Practice guidelines. The central Institutional Review Board/Ethics Committee (Advarra, Inc., 6100 Merriweather Drive, Columbia, MD 21044) approved the protocol and consent forms. All participants provided written informed consent. Eligible participants included healthy male and female adults >18 years of age. Samples from individuals who had not experienced natural SARS-CoV-2 infection (anti-nucleocapsid antibody negative and no history of clinical SARS-CoV-2 infection) were selected for analysis.

### Viral antigens

Recombinant soluble spike proteins (spike, RBD, and NTD) from Wuhan-1, B.1.351, BA.1, BA.4/5, and XBB.1.5 SARS-CoV-2 strains were expressed as described^49,50^. Recombinant proteins were produced in Expi293F cells (ThermoFisher, Cat. #A14635) by transfection of DNA using the ExpiFectamine 293 Transfection Kit (ThermoFisher, Cat. #A14524). Supernatants were harvested 4-5 days post-transfection, and recombinant proteins were purified using Strep-Tactin XT resin (IBA Lifesciences, Cat. #2-5010), then buffer exchanged into PBS and concentrated using Amicon Ultracel centrifugal filters (EMD Millipore, Cat. #UFC905096). SARS-CoV-2 (Wuhan-1 strain) S2 protein (Hexapro-SS-2W) included residues 697–1208 of the spike with an artificial signal peptide, proline substitutions at positions 817, 892, 899, 942, 986, and 987, cysteine substitutions at positions 704 and 790, a glutamate substitution at 957, tryptophan substitutions at 991 and 998, and a C-terminal foldon trimerization motif, HRV 3C cleavage site, 8× His tag and Twin-Strep-tag^51^. Soluble Hexapro- SS-2W protein was expressed and purified as described^51^ in FreeStyle 293F cells (ThermoFisher, Cat. #R79007).

For multiplexed bead-binding assays and antibody depletion assays, recombinant spike proteins were biotinylated with EZ-Link™ NHS-PEG4-Biotin (ThermoFisher, Cat. # A39259) for 2 h at 4°C and processed through Zeba spin desalting columns (ThermoFisher) to remove excess unbound biotin.

### ELISA

Purified recombinant Wuhan-1, B.1.351, BA.1, or BA.5 (2 μg/mL) spike proteins were coated onto 96-well Maxisorp plates (ThermoFisher, Cat. #439454) in 50 μL of 50 mM Na_2_CO_3_ pH 9.6 overnight at 4°C. Coating buffer was removed, and wells were washed 3 times with PBS + 0.05% Tween-20 (PBST) and blocked with PBST + 2% BSA (blocking buffer) for 1 h at room temperature. Sera were serially diluted in blocking buffer and added to the plates. Plates were incubated for 1 h at 37°C and then washed 3 times with PBST, followed by addition of 50 μL of 1:750 anti-mouse IgG-HRP (Sigma-Aldrich, Cat. #A5278) in blocking buffer. Following a 1 h incubation at 37°C, plates were washed 3 times with PBST and added with 50 μL of 1-Step Ultra TMB-ELISA (ThermoFisher, Cat. #34028). Reactions were stopped by the addition of 50 μL of 2 M sulfuric acid. Optical density measurements were taken at 490 nm using a microplate reader (BioTek). The endpoint serum dilution was calculated with curve fit analysis of optical density (OD) values for serially diluted sera with a cut-off value set to mean plus three times the standard deviation of the background signal.

### Multiplexed spike-binding assay

Recombinant biotinylated spike proteins (spike, RBD, NTD, S2) from Wuhan-1, B.1.351, BA.1, BA.5, and XBB.1.5 strains of SARS-CoV-2 were incubated with different fluorescence intensity peaks of the SPHERO Streptavidin fluorescent Yellow Particles (Spherotech, Cat. #SVF3-2552) at 20 ng per μg beads for 30 min at room temperature on an end-over-end mixer. Free biotin (Avidity, Cat. #BIO200) was added to the beads at 5 μM and incubated for 15 min at room temperature. Beads of different intensity and loaded with different spike proteins were then pooled and washed with PBS supplemented with 2% FBS, 2 mM EDTA, and 0.1% sodium azide (FACS buffer), mixed with serially diluted pre-cleared serum samples or mAbs, and incubated for 30 min at room temperature. Beads were washed twice with FACS buffer and stained with a mixture of anti-mouse IgG Alexa Fluor™ 647 (ThermoFisher, Cat. #A-21235, 1:3000) and anti-human IgG Alexa Fluor™ 647 (ThermoFisher, Cat. #A-21445, 1:3000) for 10 min at room temperature in dark. After two washes with FACS buffer, beads were resuspended in FACS buffer and acquired on flow cytometer iQue®3 (Sartorius). Population gating and analysis of fluorescence intensity were performed with iQue Forecyt software. The average of geometric mean fluorescence intensity of no-antibody control wells was defined as the background signal. Background subtracted fluorescence intensity of samples was divided by background values to obtain the fold change and interpolated with CV3-25 (spike and S2), SARS2-38 (RBD), or SARS2-57 (NTD) standard curves performed in duplicates. Interpolated values were multiplied by the sample dilution factor, and the final binding titers were shown as ng/mL equivalents of CV3-25, SARS-38, or SARS2-57.

### Antibody depletion assay

Streptavidin magnetic beads (BioLabs, Cat. #S1420S) were washed twice with PBST and incubated with recombinant biotinylated spike proteins (spike, RBD, NTD, S2) from Wuhan- 1, B.1.351, BA.1, BA.5, or XBB.1.5 strains of SARS-CoV-2 in PBS at 30 μg per mg beads for 30 min at room temperature on an end-over-end mixer. Beads were washed twice with PBS and incubated with diluted sera samples for 30 min at room temperature with agitation. Depleted sera were collected on the Kingfisher Flex extraction robot (ThermoFisher) and stored at 4°C.

### Pseudovirus neutralization assay

Vero-TMPRSS2 cells were seeded into 96 well plates in DMEM supplemented with 10% FBS without blasticidin. The next day, depleted serum samples or PBS were serially diluted with DMEM supplemented with 2% FBS and mixed with VSV pseudoviruses and incubated for 1 h at 37°C. Antibody-virus complexes were added to Vero-TMPRSS2 cell monolayers and incubated for 1 h at 37°C. Subsequently, cells were overlaid with 1% (w/v) methylcellulose in MEM. Plates were harvested 12 h (MERS), 20 h (Wuhan-1, B.1.351, BA.5, XBB.1.5, and EG.5.1), 26 h (BA.1 and SARS-CoV-1), or 30h (Pang/GD) later by removing overlays and fixed with 4% PFA in PBS for 20 min at room temperature. Plates were washed and sequentially incubated with anti-VSV nucleoprotein antibody (1:5,000 dilution, Sigma-Aldrich, Cat. #MABF2348) or anti-VSV matrix protein antibody (1:5,000 dilution, Sigma-Aldrich, Cat. #MABF2347) and subsequently with HRP-conjugated goat anti-mouse IgG (1:750 dilution, Sigma-Aldrich, Cat. #A5278) in PBS supplemented with 0.1% saponin and 0.1% bovine serum albumin. Pseudovirus-infected cell foci were visualized using TrueBlue peroxidase substrate (KPL, Cat. #5510-0050) and quantitated on an ImmunoSpot analyzer (Cellular Technology).

### Production of mAbs

To produce mAbs recombinantly, ∼15 to 30 mL ExpiCHO cell cultures were transfected using the Gibco ExpiCHO Expression System (ThermoFisher) as described by the vendor’s protocol. MAbs were purified from clarified cell culture supernatants using HiTrap MabSelect SuRe columns (Cytiva, Cat. #GE11-0034-95) on a 24-column parallel protein chromatography system (Protein BioSolutions). Purified mAbs were buffer exchanged into 1X DPBS, concentrated using Amicon Ultra 50 KDa-cutoff centrifugal filter units (Millipore Sigma), and stored at 4°C until use.

### MAb competition assay

Recombinant biotinylated XBB.1.5 spike protein was incubated with SPHERO Streptavidin fluorescent Yellow Particles (Spherotech, Cat. #SVF3- 2552) at 20 ng per μg beads for 30 min at room temperature on an end-over-end mixer. Free biotin (Avidity, Cat. #BIO200) was added to the beads at 5 μM and incubated for 15 min at room temperature. Beads were then washed with PBS supplemented with 2% FBS, 2 mM EDTA, and 0.1% NaN_3_ (FACS buffer), mixed with serially diluted serum samples, and incubated for 30 min at room temperature followed by 5 min at 4°C. Without washing, mAbs (final concentrations of 0.002 μg/mL of rCOV2-3731, rCOV2-3872, rCOV2-3967, rCOV2-3889, or rCOV2-4030, 1 μg/mL of S309, or 0.1 μg/mL of CC40.8 or CV3-25) that were pre-labeled with Alexa Fluor 647 (using Zip Alexa Fluor™ Rapid Antibody Labeling Kit (ThermoFisher, Cat. #Z11235)) were added to the beads and incubated for 30 min at room temperature in dark. After two washes with FACS buffer, beads were resuspended in FACS buffer and acquired on flow cytometer iQue®3 (Sartorius). Population gating and analysis of fluorescence intensity were performed with iQue Forecyt software. The average geometric mean fluorescence intensity (GMT) of no mAb control wells was defined as the background signal. Background-subtracted fluorescence intensity of samples was divided by the average GMT of no serum control wells to obtain % mAb binding.

### Statistical analysis

*P* values were determined using GraphPad Prism version 10. Tests (Mann-Whitney test with Bonferroni correction; Kruskal-Wallis ANOVA with Dunn’s post-test; Wilcoxon signed-rank test), number of animals, numbers of humans, median or mean values, and statistical comparison groups are indicated in the Figure legends.

## Data and reagent availability

All data supporting the findings of this study are available within the paper or its Extended Data files. Any additional information related to the study also is available from the corresponding author upon reasonable request. All reagents are available through a Material Transfer Agreement.

