## Supplementary figures and images for "Prototype mRNA vaccines imprint broadly neutralizing human serum antibodies after Omicron variant-matched boosting"

### Extended Data Figure 1

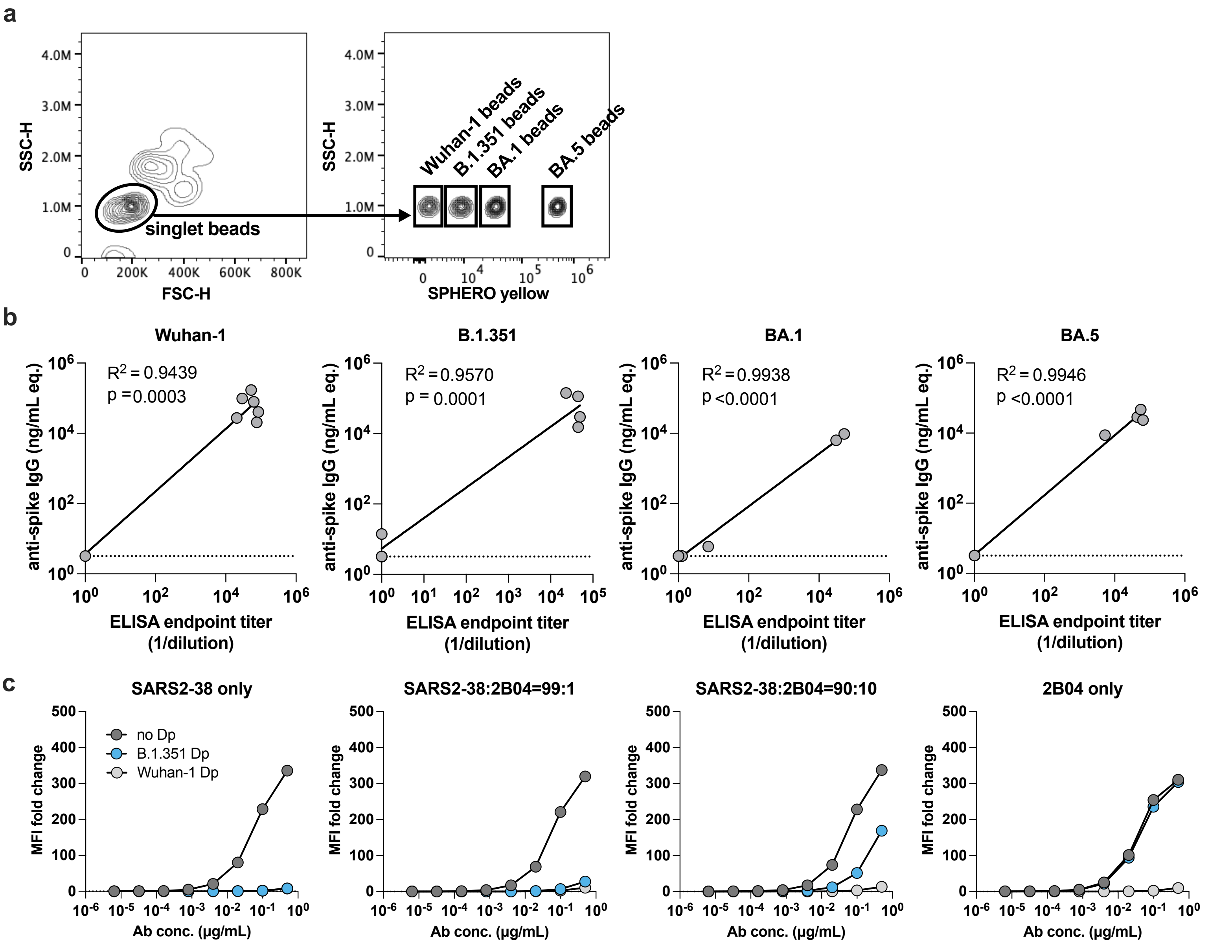


#### Extended Data Figure 1

### Extended Data Figure 2

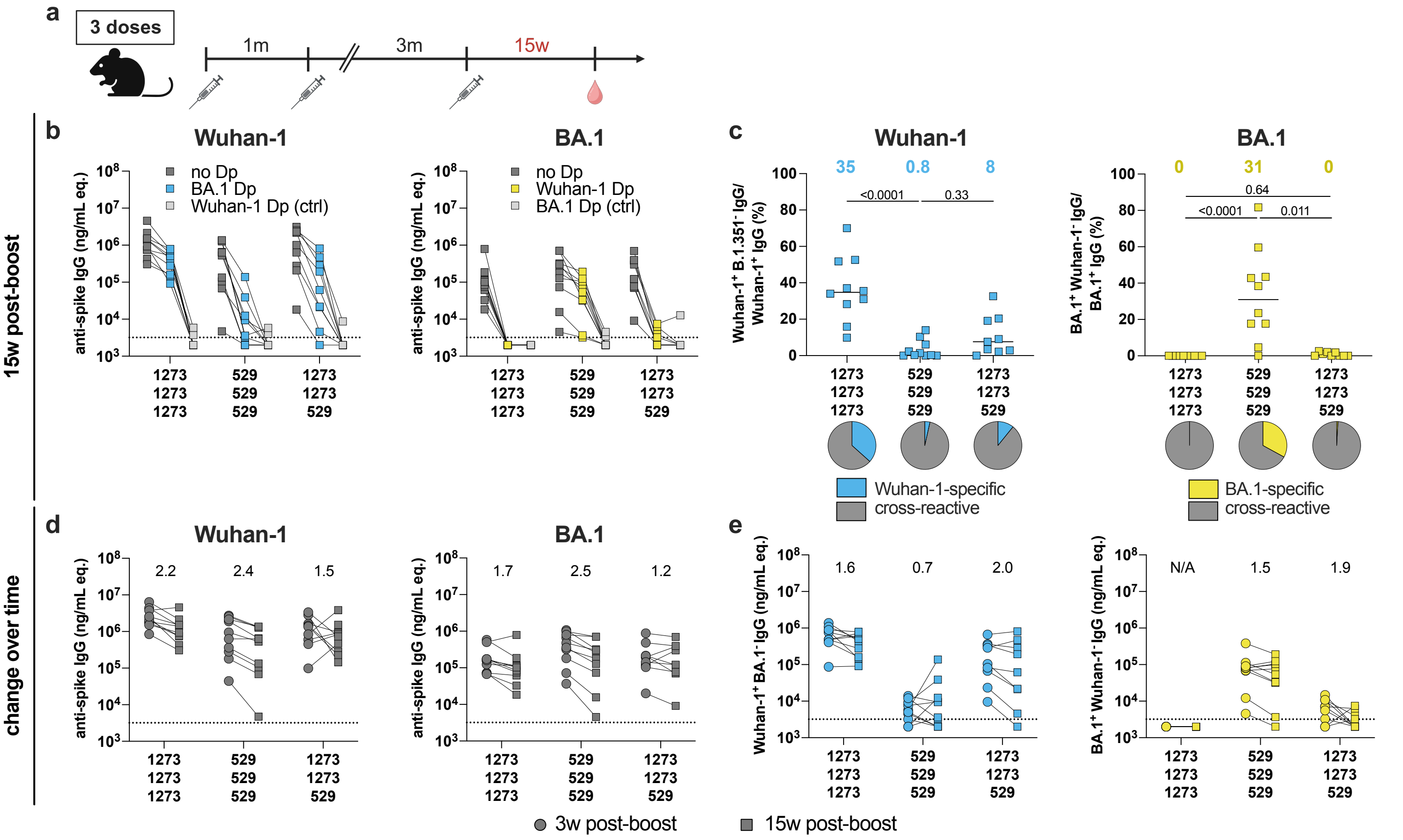


#### Extended Data Fig. 2

### Extended Data Figure 3

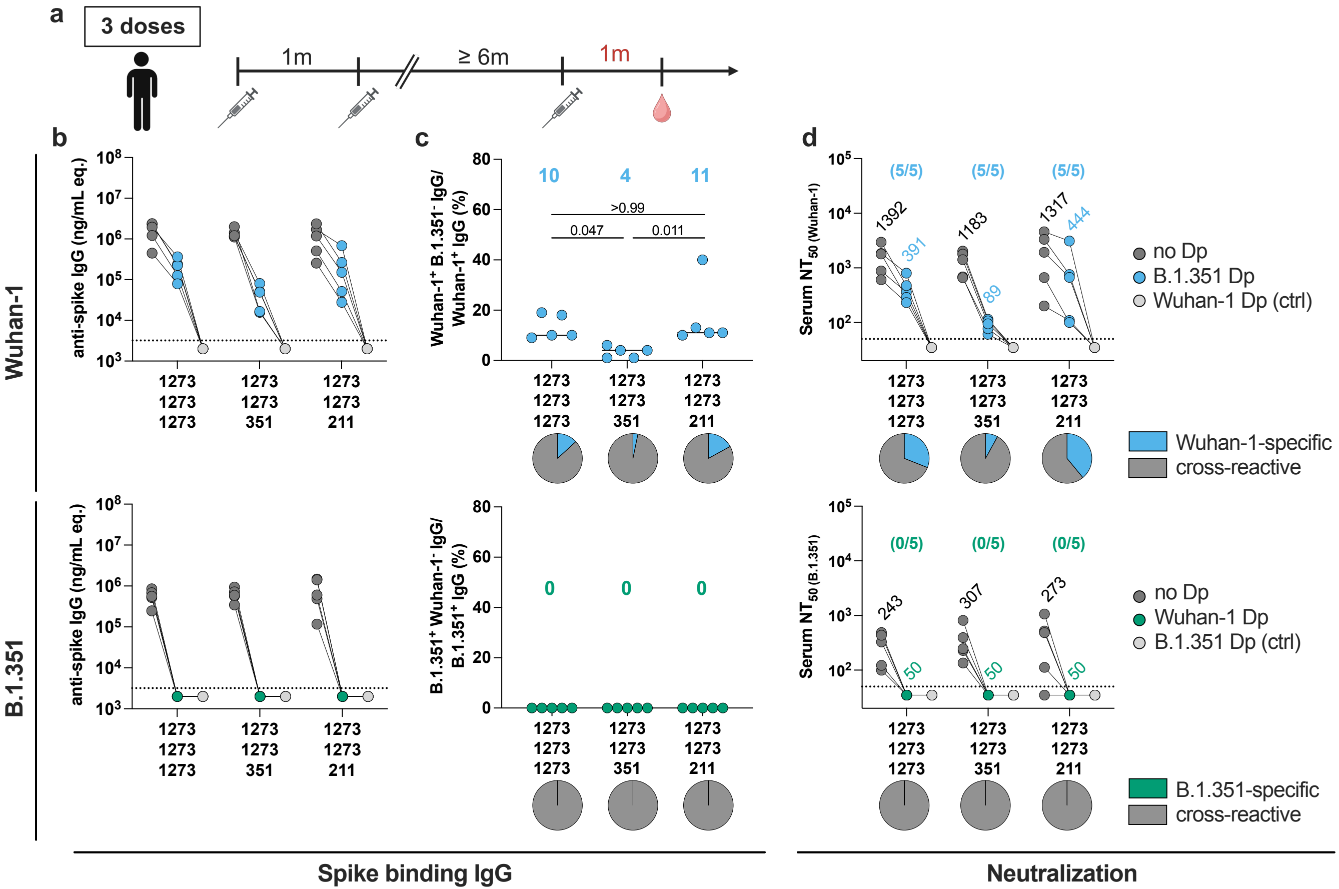


#### Extended Data Fig. 3

### Extended Data Figure 4

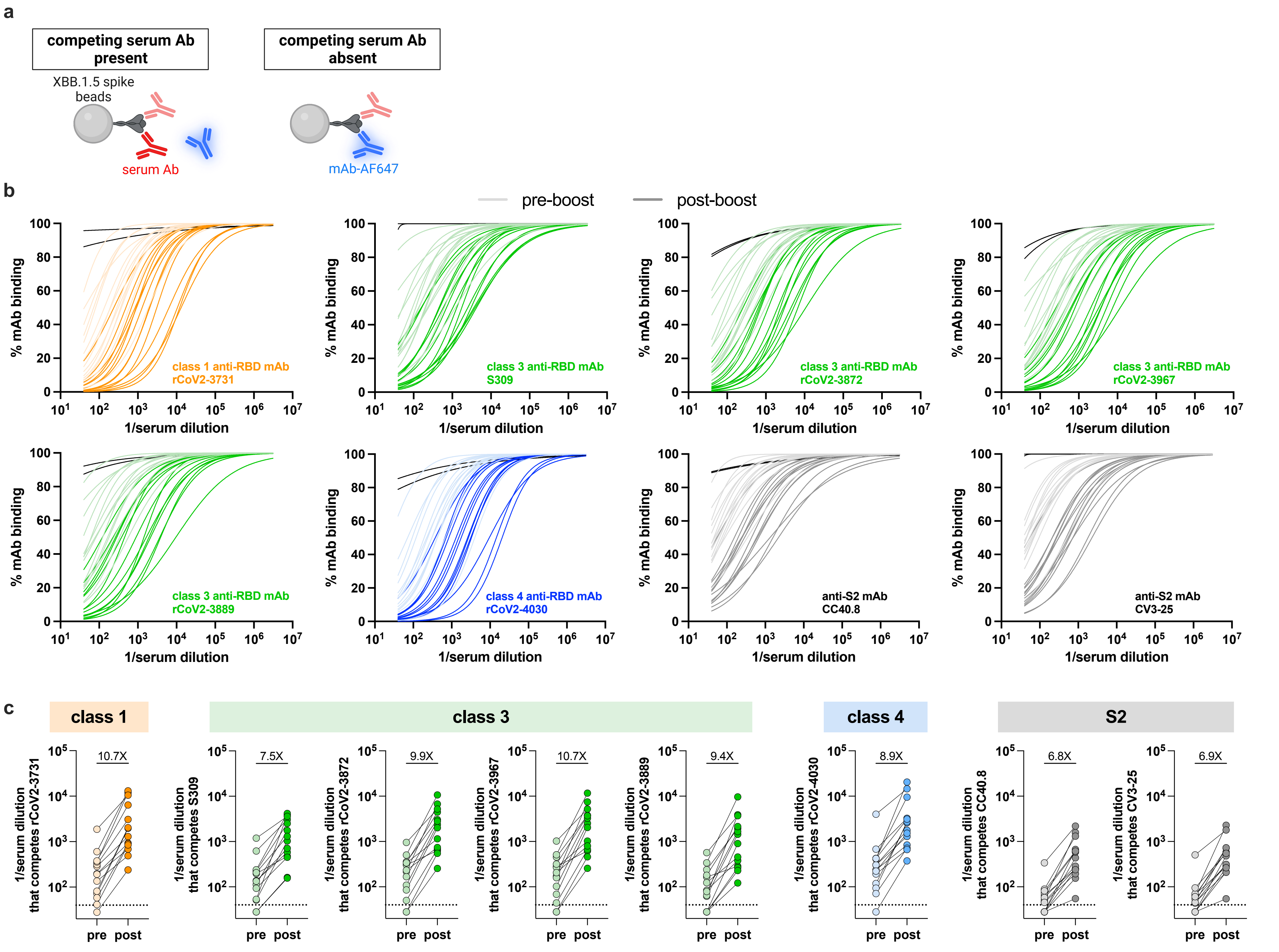


#### Extended Data Fig. 4

### Extended Data Figure 5

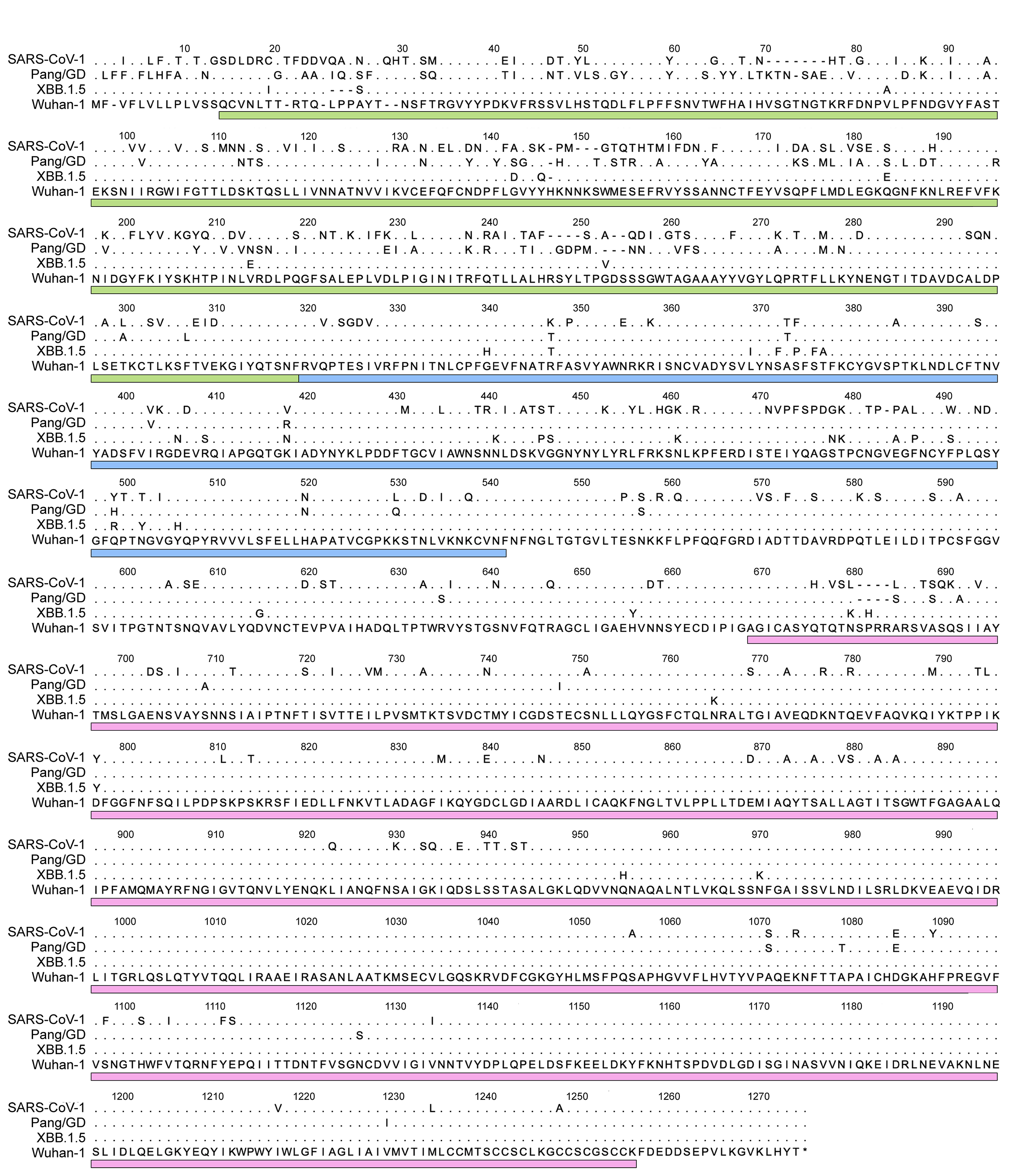

### Extended Data Figure 6

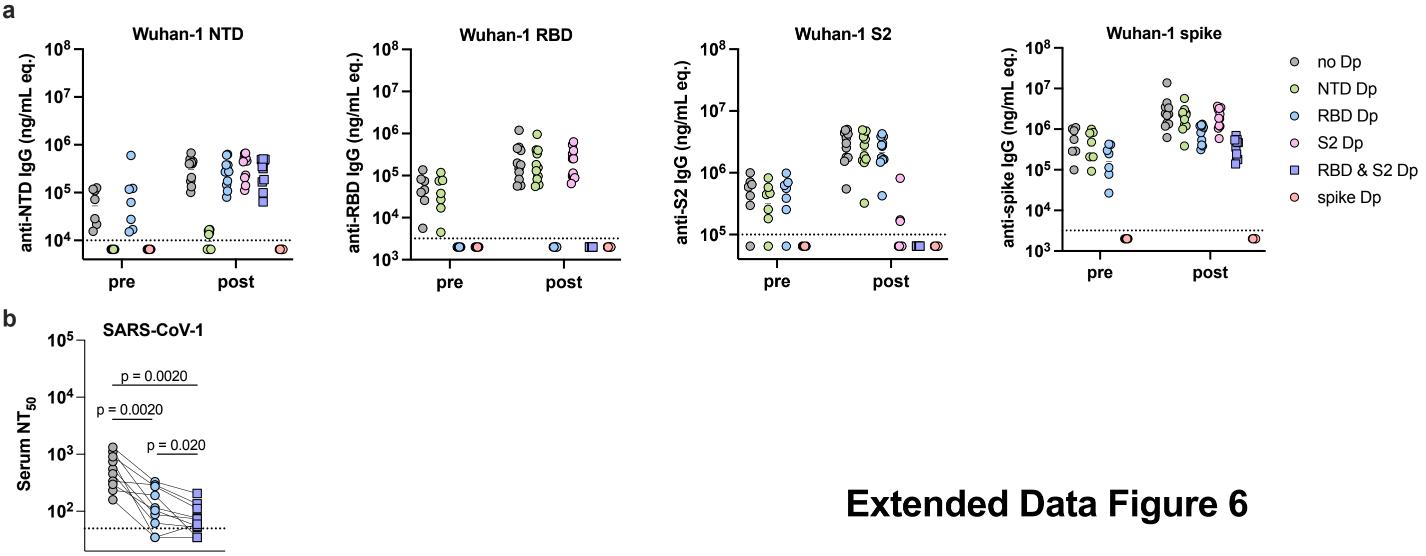


#### Extended Data Figure 6
